# Intrinsic immune receptor triggering mechanism informs rational chimeric receptor design

**DOI:** 10.64898/2026.10.24.761481

**Authors:** Songfang Wu, Hui Chen, Qi Wang, Haiyan Qiang, Xin Guan, Yong Zhang, Wei Hu, Wei Chen, Changjie Lou, Jizhong Lou

## Abstract

The T cell receptor (TCR) couple antigen recognition to T-cell activation and immune homeostasis. As an eight-subunit assembly, TCR activation operates as a highly coordinated, multi-subunit signaling event. However, which subunit serves as the primary trigger upon ligand engagement still remains unresolved. we identify CD3ε as a triggering subunit whose intracellular domain (ICD) is released from the plasma membrane during TCR activation. Elevated membrane tension facilitates this transition from a membrane-proximal to a membrane-distal conformation. The released ICD forms condensates through phase separation with free or co-receptor-associated Lck. These condensates recruit additional CD3ε ICDs from neighboring TCRs, triggering resting receptors and promoting further recruitment of TCRs and Lck to amplify proximal signaling. The CD3ε ICD alone support robust T cell activation independently of its own ITAM. We observe a similar triggering mechanism in several activating receptors in T and nature killer (NK) cell. These findings support a unifying model in which designated subunits initiate multi-subunit immune receptor activation through phase separation with key kinases. Guided by this model, we engineered triggering molecules (TRIG) that lack canonical intracellular signaling modules. These TRIG molecules recruit endogenous immune receptors in a ligand-dependent manner and efficiently activate both T cells and NK cells. This mechanism provides a basis for designing chimeric receptors that activate immune cells through endogenous receptor signaling.

## Introduction

T cell activation depends on the recognition of antigenic peptides presented by major histocompatibility complex (MHC) by T cell receptor (TCR). The canonical αβ TCR complex comprises two highly variable chains (α and β) assembled with three signaling dimers: CD3εδ, CD3εγ, and CD3ζζ (Rudolph et al., 2006). Extracellular TCR–pMHC binding kinetics are transduced into intracellular signaling events, prompting the phosphorylation of immunoreceptor tyrosine-based activation motifs (ITAMs) across the CD3 subunits to initiate downstream cascades (Courtney et al., 2018). Although structural biology has elucidated cryo-EM structures of the TCR–pMHC (Dong et al., 2019; Sušac et al., 2022), the precise physical mechanism that triggers TCR activation remains elusive.

Current paradigms explain TCR activation through three primary, non-mutually exclusive models: receptor aggregation (clustering), conformational changes (driven by ligand binding or mechanical force), and kinetic segregation (size-dependent exclusion of large phosphatases like CD45) (Travaglino et al., 2026; van der Merwe and Dushek, 2011). Upon antigen engagement, the T-cell and target-cell membranes form a tight junction that physically excludes CD45 (Courtney et al., 2018). Simultaneously, mechanical tension transmitted across the extracellular TCR induces structural rearrangements that shift the receptor into a pre-activated state (Wu et al., 2019). In this state, the TCR undergoes phase separation and becomes phosphorylated by free or co-receptor-associated Lck, promoting receptor micro-clustering (Chen et al., 2023). Phosphorylated TCR clusters then recruit and activate Zap70, propagating signals to downstream scaffolds such as LAT (Su et al., 2016). However, dissecting the precise interplay between mechanical tension, CD3 conformational changes, and kinetic segregation has proven technically challenging.

Structural analyses reveal that the extracellular domains of the TCR undergo minimal conformational change upon pMHC binding (Natarajan et al., 2024; Xu et al., 2020). In contrast, the intracellular domains (ICDs) of the CD3ε subunit and CD3ζ exhibit marked conformational rearrangements upon pMHC engagement (Alarcon and Schamel, 2025; Lanz et al., 2021; Mariuzza et al., 2020; Minguet et al., 2007). In the resting TCR, the ICDs of CD3ε and CD3ζ associate with negatively charged phospholipids in the plasma membrane through their basic-rich sequences (BRS), maintaining a membrane-proximal, concealed conformation. Upon TCR ligation by pMHC, these domains detach from the membrane and become exposed (Aivazian and Stern, 2000; Gil et al., 2002; Gil et al., 2005; Risueño et al., 2005; Risueño et al., 2008). Notably, the phosphorylation rates of CD3ε and CD3ζ during TCR triggering are not significantly different, although the overall phosphorylation level of CD3ζ is clearly higher than that of CD3ε (Chen et al., 2025; Wu et al., 2020). When individual phosphorylation sites are examined, both CD3ε and CD3ζ exhibit considerable heterogeneity, with membrane-proximal tyrosine being preferentially phosphorylated (Wu et al., 2020). However, the signaling hierarchy of CD3ε and CD3ζ and their respective roles throughout the TCR triggering process remain poorly defined. Furthermore, it is unclear whether the ICDs of CD3δ and CD3γ undergo similar conformational changes upon pMHC binding.

Clustering, a common phenomenon during receptor triggering, plays a crucial role in receptor signaling (Grakoui et al., 1999). TCRs are evenly distributed on naïve T cells but form nano-clusters in antigen-experienced T cells. In the early stage of T cell activation, pMHC ligated TCR molecules condensate with Lck kinase to form micro-clusters, which mediate CD3 phosphorylation and downstream signal transduction (Demetriou et al., 2020; Hashimoto-Tane et al., 2016). Subsequently, TCR clusters are transported to the center of cell-cell contact region and form the central supramolecular activation cluster (cSMAC), surrounded by other receptor clusters, giving rise to the typical bull’s eye structure known as the immunological synapse (IS) (Hashimoto-Tane et al., 2011). CD3ε condensation with Lck has been identified as a unique property of the TCR and plays a key role in TCR clustering and IS assembly (Chen et al., 2023). CD3ε/Lck condensation unleashes the kinase activity of Lck and promotes CD3 phosphorylation, in turn, phosphorylation further enhances CD3ε/Lck condensation. This self-promoting phase separation system drives proximal signal amplification and helps TCR surpass the activation threshold, potentially representing a checkpoint event in TCR triggering.

Unlike the TCR/CD3 complex, chimeric antigen receptors (CARs) are designed by incorporating an antigen recognition domain, typically a single-chain variable fragment (scFv), followed by a hinge region, a transmembrane region, and an intracellular signaling sequence. The intracellular portion contains one (the second-generation CARs) or two (the third-generation CARs) costimulatory domains, fused to the intracellular region of CD3ζ (Wu et al., 2022). This modular design allows extensive variation in antigen recognition and signaling, enabling CARs to be widely used in therapy of cancer and autoimmune diseases (Ellebrecht et al., 2016; Hunter et al., 2025; Jin et al., 2021; Wang et al., 2024). Although CARs have achieved remarkable success in treating certain hematological malignancies, their clinical efficacy in other hematological malignancies and solid tumors remains limited. One important reason may be the insufficient diversity of ITAMs in current CAR designs. Recent efforts have sought to overcome these limitations by modulating the types and diversity of ITAMs. Studies have shown that ITAM diversity is critical for T cell proliferation (Guy et al., 2013), and that a greater variety of ITAMs can increase the fraction of cells that become activated, rather than simply enhancing signal strength in individual cells (Mayya and Dustin, 2016). In a direct comparison, TCR and CAR were co-expressed in single cells, although TCR-induced signal was slower and more gradual than CAR signaling, it was significantly more sustained (Davenport et al., 2018).

Capitalizing on the advantages of TCR signaling, the TRuC-T design harnesses the complete endogenous TCR complex by fusing a single-chain antibody (ScFv) to the extracellular region of CD3ε. This approach achieves potent tumor killing with lower cytokine secretion, thereby reducing the risk of cytokine release syndrome (Baeuerle et al., 2019). The signal events induced by TRuC-T are consistent with the activation of the entire TCR of the cell, for example, it causes the phosphorylation of CD3ε and CD3ζ, but this is not observed in traditional CARs. Another alternative to traditional CARs is the STAR-T/HIT design, which replaces the variable regions of the TCRα and β receptors with the heavy and light chains of an ScFv, thereby also engaging the endogenous TCRs. These designs retain the overall architecture and ITAM diversity of endogenous TCRs, endowing T cells with improved memory and durable therapeutic efficacy (Liu et al., 2021; Mansilla-Soto et al., 2022; Woessner et al., 2025). Compared with TCR-engineered T cells (TCR-T), they offer great target flexibility and therefore hold considerable potential for the treatment of tumors and autoimmune diseases.

In this study, we found CD3ε-ICD transitions from a membrane-proximal conformation in the resting state to a membrane-distal conformation during TCR triggering, a process facilitated by increased membrane tension. Once released from the plasma membrane, CD3ε-ICD can bind to free or co-receptor-associated Lck and undergo condensation. The resulting condensates can further recruit other CD3 subunits, drawing resting TCR and Lck into the condensates and thereby achieving proximal signal amplification. Importantly, CD3ε-ICD can mediate T cell activation independently of its own ITAM and act as a trigger for the TCR. Unlike the direct signal transmission by CD3ζ in conventional CAR-T cells, TRIG-T can recruit endogenous TCR molecules, generating a signal that resembles endogenous TCR engagement by pMHC and thus endowing T cells with tumor-killing capacity. We also found that certain NK receptors possess trigger subunits that, like CD3ε, undergo phase separation with Lck. Additionally, we designed similar trigger molecules in silico, these TRIG molecules trigger T and NK cell activation in manner dependent on their phase separation with Lck. TRIG mediates signaling recapitulates endogenous receptor activation, offering a new potential strategy for immune cell engineering.

## Results

### CD3ε chain may serve as a trigger within TCR complex

It has been reported that the cytoplasmic domain of CD3ε and CD3ζ undergo conformational changes upon TCR engagement with agonist pMHC. To dissect TCR triggering mechanism, we first examined whether the intracellular domain (ICD) of CD3δ and CD3γ can associate with the plasma membrane in T cells using a fluorescence resonance energy transfer (FRET) assay. We generated four chimeric antigen receptor (CAR) constructs based on the ICDs of the CD3ε, CD3ζ, CD3δ, or CD3γ. Each CAR contains an N terminus CD8 signal peptide, a human CD19-targeted single-chain fragment variable (FMC63), the CD8α hinge and transmembrane domains, and the respective CD3 ICD fused to mTFP1. These four CD3s-CAR Jurkat cells were labeled with the lipid dye R18 and stimulated on glass surfaces coated with human CD19 protein or antibodies (**Figure S1A**). This approach allowed us to assess the proximity between the CD3 ICDs and the plasma membrane by measuring the FRET efficiency between mTFP1 and R18 (**Figure S1B**). In the resting state, CD3δ-ICD and CD3γ-ICD displayed negligible FRET efficiency (**Figure S1C**), in contrast to CD3ε-ICD and CD3ζ-ICD, which have been reported to bind the plasma membrane. However, the membrane association of CD3ε-ICD and CD3ζ-ICD was largely abrogated upon stimulation with CD19 and ScFv-targeted antibody (**Figure S1D**). The results also revealed that CD28-signaling and endogenous TCR-signaling exerted minimal effects on the interaction of CD3ε-CAR with the plasma membrane (**Figure S1D**). Our findings suggest that both CD3ε and CD3ζ bind membrane lipids in the resting state and are released upon TCR triggering, thus potentially acting as the first responders in the TCR triggering process.

Liquid-liquid phase separation between CD3ε and Lck have been reported as an initial response during TCR triggering, however, the relationship between CD3ε membrane binding and its phase separation with Lck remain unknown. To address this question, we first coated His-tagged CD3ε-ICD on a supported lipid bilayer (SLB) to allow its interaction with lipids, and then added His-tagged Lck_UD-SH3-SH2_. We found that both CD3ε-ICD and Lck_UD-SH3-SH2_ exhibited a uniform distribution on the SLB (**Figure 1A**). This result indicates that membrane binding may shield a key motif of CD3ε and thereby provent its phase separation with Lck. To validate the system, we subsequently added soluble CD3ε to the solution to trigger Lck_UD-SH3-SH2_ condensation, and observed that Lck_UD-SH3-SH2_ formed distinct clusters on SLB. Surprisingly, we also found that His-tagged CD3ε-ICD was enriched in these Lck_UD-SH3-SH2_/CD3ε clusters, and a similar result was observed when soluble CD3ε-pY1 was used (**Figure 1A**). We further examined the other three CD3 subunits and found that only CD3ζ could induce slight condensation of Lck_UD-SH3-SH2_ together with His-tagged CD3ε (**Figure 1B**). Moreover, the Lck_UD-SH3-SH2_/CD3ε clusters could recruit other CD3 subunit that had been pre-incubated with the SLB (**Figure 1C**). Collectively, these results suggest a triggering role for CD3ε within the TCR complex, whereby it induce Lck clustering and subsequently recruits other CD3 ICDs in the resting state.

**Figure 1.**
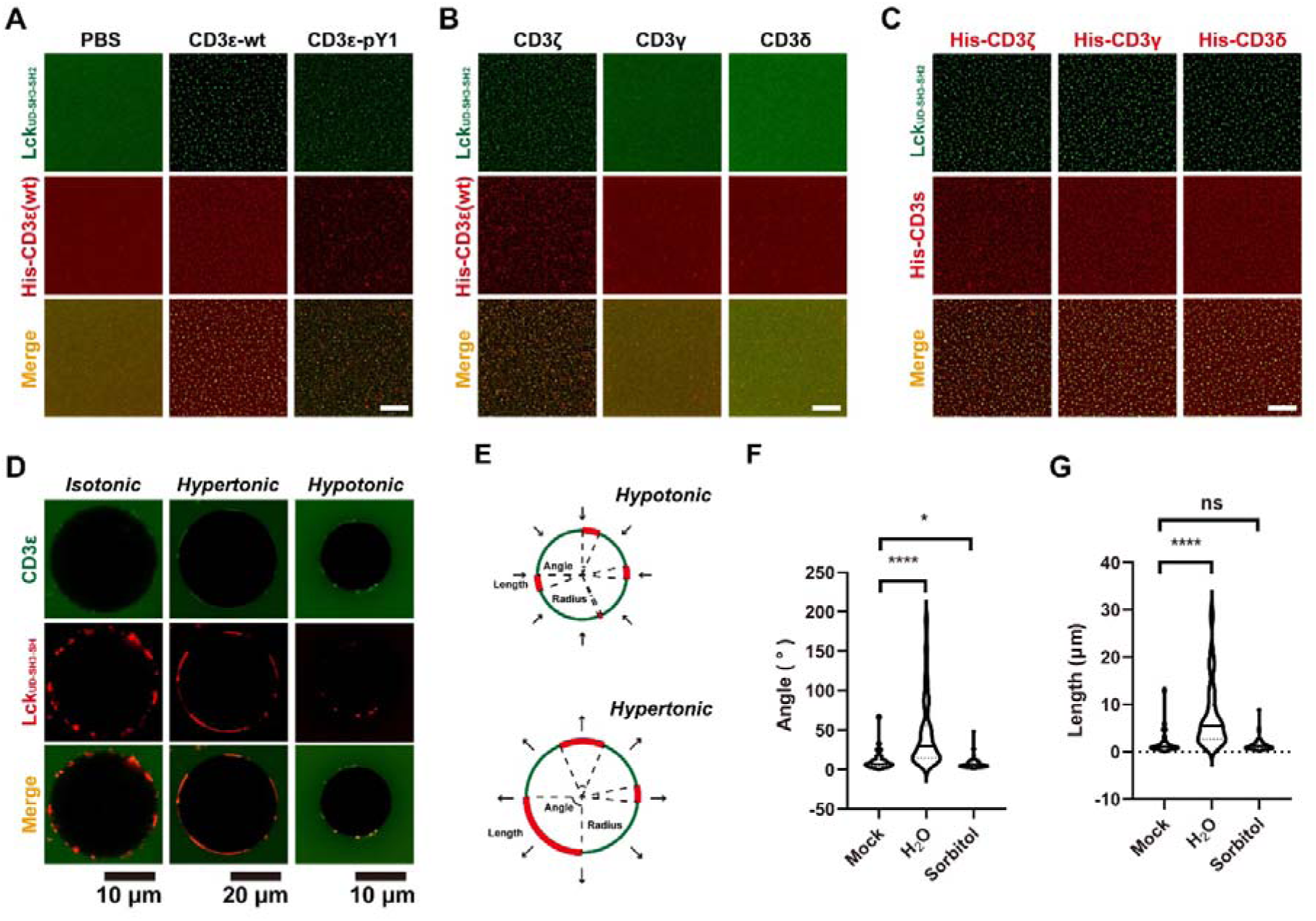
Membrane tension facilitates the phase separation between membrane-bound CD3ε and Lck kinase. **A-C**. Typical images of the phase separation of Lck_UD-SH3-SH2_ (labelled with AF-488) and membrane-bound CD3s (His-CD3ε in **A** and **B**, His-CD3ζ, His-CD3γ and His-CD3δ in **C**, all labelled with AF-647) on SLB. Phase separation was triggered with soluble CD3 supplement (PBS mock, CD3ε and CD3ε-pY1 in **A**, CD3ζ, CD3γ and CD3δ in **B**, CD3ε in **C**), Scale bar: 10 μm **D-G**. Phase separation of Lck_UD-SH3-SH2_ (labelled with AF-647) and CD3ε (labelled with AF-488) on GUV under different osmotic conditions. Typical images and schematic diagram were shown in **D** and **E**, the statistics of the angle and length were shown in **F** and **G**. Data were shown as Violin plus Medium and Quartiles (n=31 for Mock; n=86 for H_2_O treated; n=85 for Sorbitol treated; Unpaired t test; ns, *P* > 0.05; *, *P* < 0.05; ****, *P* < 0.0001).

TCR has long been recognized as a mechanical force sensor that binds non-self/agonist pMHC via catch bond and self/antagonist pMHC with slip bond. The force generated by extracellular ligand binding can be transmitted through the transmembrane domain to the membrane and its intracellular domains. Our result, together with previous studies, have revealed that ligand binding facilitates the release of CD3ε-ICD from the membrane, suggesting that membrane tension may serve as an important regulator in this process. To characterize how membrane tension regulates the conformational change of CD3ε, we designed a GUV-based condensation system. We sequentially ligated His-tagged CD3ε and Lck_UD-SH3-SH2_ onto GUV surface, and then used PBS, H_2_O or sorbitol to individually adjust the membrane tension of GUVs. CD3ε and Lck_UD-SH3-SH2_ formed clusters under all three conditions (**Figure 1D**). We next quantified the size of the clusters by measuring their angel and length on the GUV surface (**Figure 1E**). Our results showed that hypertonic state induced by H_2_O promoting the condensation of CD3ε/Lck_UD-SH3-SH2_ on the surface (**Figure 1F and G**). To further exclude ionic effects, we established an SLB-based membrane tension system in which membrane tension was generated by cholesterol depletion from pre-coated SLB. His-tagged CD3ε and Lck_UD-SH3-SH2_ were sequentially coated onto the SLB, after which MβCD was applied to remove cholesterol and generate membrane tension, and soluble CD3ε was then added to induce phase separation (**Figure S1E**). The results demonstrate that membrane tension (cholesterol depletion) significantly enhanced CD3ε/Lck_UD-SH3-SH2_ condensation (**Figure S1F and G**).

Together, our results suggest that the CD3ε subunit may serve as a triggering element within the TCR complex. Upon TCR binding to agonist pMHC, mechanical force is transduced to the membrane and CD3ε-ICD, promoting the release of CD3ε-ICD from the plasma membrane. The liberated CD3ε-ICD then rapidly forms condensates with co-receptor associated or free Lck, and these CD3ε/Lck condensates can further recruit other CD3 subunits, thereby facilitating the phosphorylation of CD3 ITAMs.

### Extracellular phase separation-mediated CD3ε ICD crowding promotes but does not trigger T cell activation

Soluble antibodies (anti-CD3 and anti-CD28) are commonly used as TCR activators in immunological studies, as they can initiate T cell activation by induce TCR clustering. Given that agonist pMHC triggers the TCR through a force induce phase separation mechanism, we were wondering whether phase separation itself could bypass mechanical force to initiate TCR activation. To test this, we employed a previously reported membrane-tethered LLPS systems to manipulate receptors clustering (Li et al., 2022). The extracellular domain of CD3s-CAR was replaced with PDZ, and an additional mGFP was fused to the C terminus (PDZ-CDε or CD3ε-TRIG). The LLPS system comprises a multivalent scaffold consisting of SUMO5 and SIM6-PV, in which PV is a C-terminal hexapeptide (KKETPV) that binds to PDZ (**Figure S2A**). We purified SUMO5 and SIM6-PV proteins and confirmed their LLPS capacity in vitro, testing different concentrations of SUMO5 (**Figure S2B**). We further validated this phase separation system in Jurkat cells expressing PDZ-CDε-GFP (CD3ε-TRIG). The results showed that SUMO5 and SIM6-PV formed condensates that localized tn the surface of Jurkat cells, and 5 μM SIM6-PV combined with 2 μM SUMO5 was sufficient to induce detectable condensates (**Figure S2C**). PDZ-CD3ε (CD3ε-TRIG) receptors were clearly enriched within these SUMO5/SIM6-PV condensates (**Figure 2A and B**), whereas neither SUMO5 nor SIM6-PV alone triggered the clustering of PDZ-CD3ε (CD3ε-TRIG) (**Figure S2D**).

**Figure 2.**
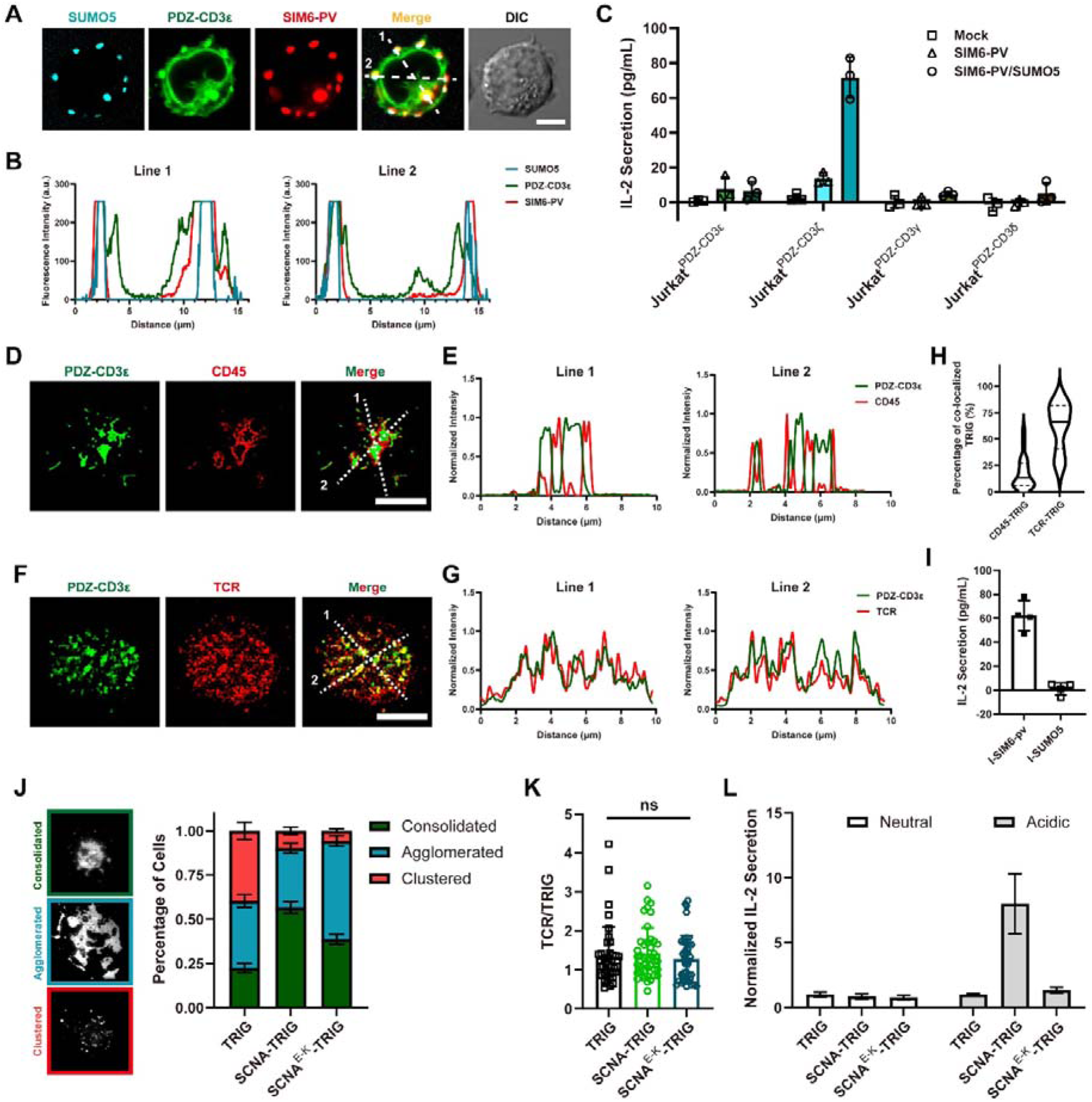
Phase Separation of the extracellular domain of PDZ-CD3ε promotes rather than directly triggers T Cell Signaling. **A** and **B**. Typical images (**A**) and co-localization analysis (**B**) of SUMO5/SIM6-PV induced clustering of PDZ-CD3ε (CD3ε-TRIG), Scale bar: 10 μm; **C**. SUMO5/SIM6-PV phase separation induced IL-2 secretion of PDZ-CD3s engineered Jurkat cells, Data were shown as Mean±SD, n=3; **D-H**. Co-localization analysis of PDZ-CD3ε (CD3ε-TRIG) with CD45 (**D**, **E** and **H**) and TCR (**F**, **G** and **H**), the percentage of TRIG molecules colocalized with CD45 (n=29) or TCR (n=40) were calculated separately (H). PDZ-CD3ε (CD3ε-TRIG) engineered Jurkat cells were stimulated with SLB-immobilized SIM6-PV, cells were fixed and stained with APC ligated CD45 or TCR antibody, Scale bar: 10μm; **I.** IL-2 secretion of PDZ-CD3ε (CD3ε-TRIG) engineered Jurkat cells stimulated with surface immobilized SIM6-PV or SUMO5, Data were shown as Mean±SD, n=4; **J.** TIRF-SIM imaging of TRIG engineered Jurkat cells, the distribution of TRIG molecules was category into three patterns (Consolidated, Agglomerated and Clustered), Left: typical images, right: percentages of three patterns, data were shown as Mean±SEM, n=10; **K.** Histogram of the TCR/TRIG ratio was calculated from **J**, data were shown as Mean±SD (TRIG: n=40; SCNA-TRIG: n=41; SCNA^E-K^-TRIG: n=41; Unpaired t test; ns, *P* > 0.05); **L.** Normalized IL-2 secretion of TRIG engineered Jurkat cells, data were shown as Mean±SD, n=3.

We next examined whether LLPS-induced condensation of PDZ-CD3 receptors could mediate Jurkat cells activation. Surprisingly, only PDZ-CD3ζ Jurkat cells produced detectable IL-2 (**Figure 2C**). CD69 expression also indicated that PDZ-CD3ε (CD3ε-TRIG) Jurkat cells could not be activated by SUMO5/SIM6-PV phase separation (**Figure S2E**). The IL-2 secretion by PDZ-CD3ε (CD3ε-TRIG) Jurkat cells stimulated with surface-ligated CD3 antibody suggested that mechanical force is necessary for CD3ε triggering (**Figure S2F**). CD45 exclusion has been established as an essential factor during T cell activation. Our previous results showed that CD3ε-ICD/Lck condensates can recruit CD45 in vitro, thus, SUMO5/SIM6-PV induced PDZ-CD3ε (CD3ε-TRIG) clusters may also recruit CD45 and subsequently attenuate downstream signaling. We therefore examined CD45 localization in PDZ-CD3ε (CD3ε-TRIG) Jurkat cells stimulated with SUMO5/SIM6-PV in solution. The results clearly showed that CD45 was enriched in PDZ-CD3ε (CD3ε-TRIG) clusters (**Figure S2G**), whereas CD45 was excluded from PDZ-CD3ε (CD3ε-TRIG) clusters when stimulated with SLB ligated SIM_6_-PV (**Figure S2H**). Collectively, these results indicate that engineered phase separation at the extracellular domain can partially trigger CD3ε condensation, but failed to transduce signaling due to impaired CD45 exclusion.

To further test this hypothesis, SIM6-PV was immobilized on an SLB surface to stimulate PDZ-CD3ε (CD3ε-TRIG) Jurkat cells, and the distribution of CD45 and TCR were quantified by TIRF-SIM imaging. We found that PDZ-CD3ε (CD3ε-TRIG) formed clusters upon immobilized SIM6-PV stimulation, and CD45 was excluded from these clusters while endogenous TCR was enriched within them (**Figure 2D-H**). IL-2 secretion results further confirmed that immobilized SIM_6_-PV could mediate the activation of PDZ-CD3ε (CD3ε-TRIG) Jurkat cells (**Figure 2I**). Together, these findings indicate that extracellular phase separation can partially elicit a CD3ε response, but failed to transduce downstream signaling due to impaired CD45 exclusion.

Given that phase separation alone is insufficient to fully activate CD3ε signaling, we next asked whether extracellular phase separation could be harnessed to promote CD3ε mediated T cell activation. Recent studies have shown that α-Synuclein (SCNA)can undergo liquid-liquid phase separation (LLPS) during its fibrillization in Parkinson’s disease pathogenesis, and that a low pH environment promotes its aggregation (Matsuo et al., 2024; Ray et al., 2020). We inserted the SCNA NAC domain into the extracellular region of the CD3ε-TRIG construct between the FMC63 and the CD8 hinge. Additionally, we generated an E46K mutation in the SCNA NAC domain, which attenuates the LLPS propensity of SCNA. Immunological synapse (IS) formation was used to quantify receptor clustering, and the results showed that TRIG receptors incorporating the SCNA NAC domain formed more compact clusters (consolidated), indicative of stronger receptors clustering (**Figure 2J**). However, the recruitment of endogenous TCR in SCNA-TRIG cells was comparable to that TRIG and SCNA^E46K^-TRIG cells (**Figure 2K**). We also evaluated the function of these SCNA NAC domain-incorporating TRIG Jurkat cells in neutral or acidic culture medium, and found that low PH enhanced T cell activation, as evidenced by increased IL-2 secretion (**Figure 2L**).

Our results demonstrate that incorporation of an extracellular phase separation module into a CD3ε ICD-based CAR can induce spatially confined crowding of CD3ε ICD under specific conditions. However, this apparent clustering is insufficient to fully trigger CD3ε/Lck phase separation. This may be attributed to incomplete release of CD3ε ICD from the plasma membrane (resulting in weak CD3ε-Lck interaction) and impaired exclusion of CD45 (which inhibits phosphorylation within CD3ε/Lck condensates). Notably, the introduction of a conditionally activated extracellular phase separation module can further amplify and propagate signals downstream of CD3ε triggering, offering a potential strategy to exploit the local tumor microenvironment for enhancing T cell activation.

### Endogenous TCR complex is indispensable for the activation of CD3ε-TRIGgering-T cells in regardless of CD3ε ITAM motif

To determine whether endogenous TCR is necessary for CD3ε-TRIG mediated T cell activation in CD3ε-TRIG-T cells, we performed assays to elucidate the detailed mechanism of CD3ε -TRIG. Jurkat T cell or J76 T cell were transduced with CD3ε-TRIG, the surface expression of CD3ε-TRIG was verified by anti-Mouse Fab antibody staining, and both CD3ε-TRIG-T cells showed similar surface expression (**Figure S3A**). Successfully transduced CD3ε-TRIG-T cells were sorted and cocultured with the Raji B cells at effector-to-target ratios of 1:0, 1:1 or 1:5. After 20 hours of co-culture, the supernatant was collected for cytokine detection. CD3ε-TRIG-Jurkat expressed IL-2 in a dose dependent manner, however the CD3ε-TRIG-J76 secreted only very weak IL-2 compared with CD3ε-TRIG-Jurkat (**Figure 3A**). Owing to the deficiency of TCR complex in J76 cells, we speculated the TCR complex is important in CD3Ε-TRIG induced T cell activation. To further substantiate this phenomenon, we then used the first-generation CD3Z-CAR, which is confirmed to activate the T cell in an TCR-independent manner. As expected, CD3Z-CAR induced significant IL-2 production in both wild type Jurkat and JRT3 cells (TCRβ-deficient), whereas CD3ε-TRIG only produced comparable IL-2 only in wild type Jurkat but not JRT3 cells (**Figure 3B**). We then mutated the two tyrosine residues of the CD3ε ITAM, individually or both simultaneously, to phenylalanine to generate non-phosphorylation mutants. As expected, we found that both inactivation of the two tyrosine residues showed stronger IL-2 expression than wild type CD3ε, while single tyrosine mutation could not induce significant IL-2 expression (**Figure 3C**). Thus, these data confirmed our hypothesis that CD3ε-TRIGgering the T cell activation via the recruitment of endogenous TCR complex instead of through the phosphorylation of its own tyrosine in ITAM motif, which was important for CD3ζ induced T cell activation. Moreover, when the 1G4 TCR was reconstituted in CD3ε-TRIG-J76 T cells, CD3ε-TRIG-induced T cell activation was significant enhanced (**Figure S3B and C**). We then performed the similar assays for the CD3δ-CAR and CD3γ-CAR and found that neither CD3δ-CAR nor CD3γ-CAR show comparable TCR dependence (**Figure S3D and E**). To generalize the finding, we substituted the transmembrane domain to CD28 TM, and similar phenomenon was detected (**Figure S3E and F**). To confirm the versatility of our CAR approach, we similarly engineered the CD3ε with a human BCMA antigen targeting CAR. We noted that the CD3ε-CAR showed comparable IL-2 expression to that of CD3Z-CAR and BBZ-CAR (**Figure S3G and H**). We also compared the four CD3 subunits-based CAR-Ts expression of CD69 as a marker of T cell activation when cocultured with the Raji B cells (**Figure S3I**).

**Figure 3.**
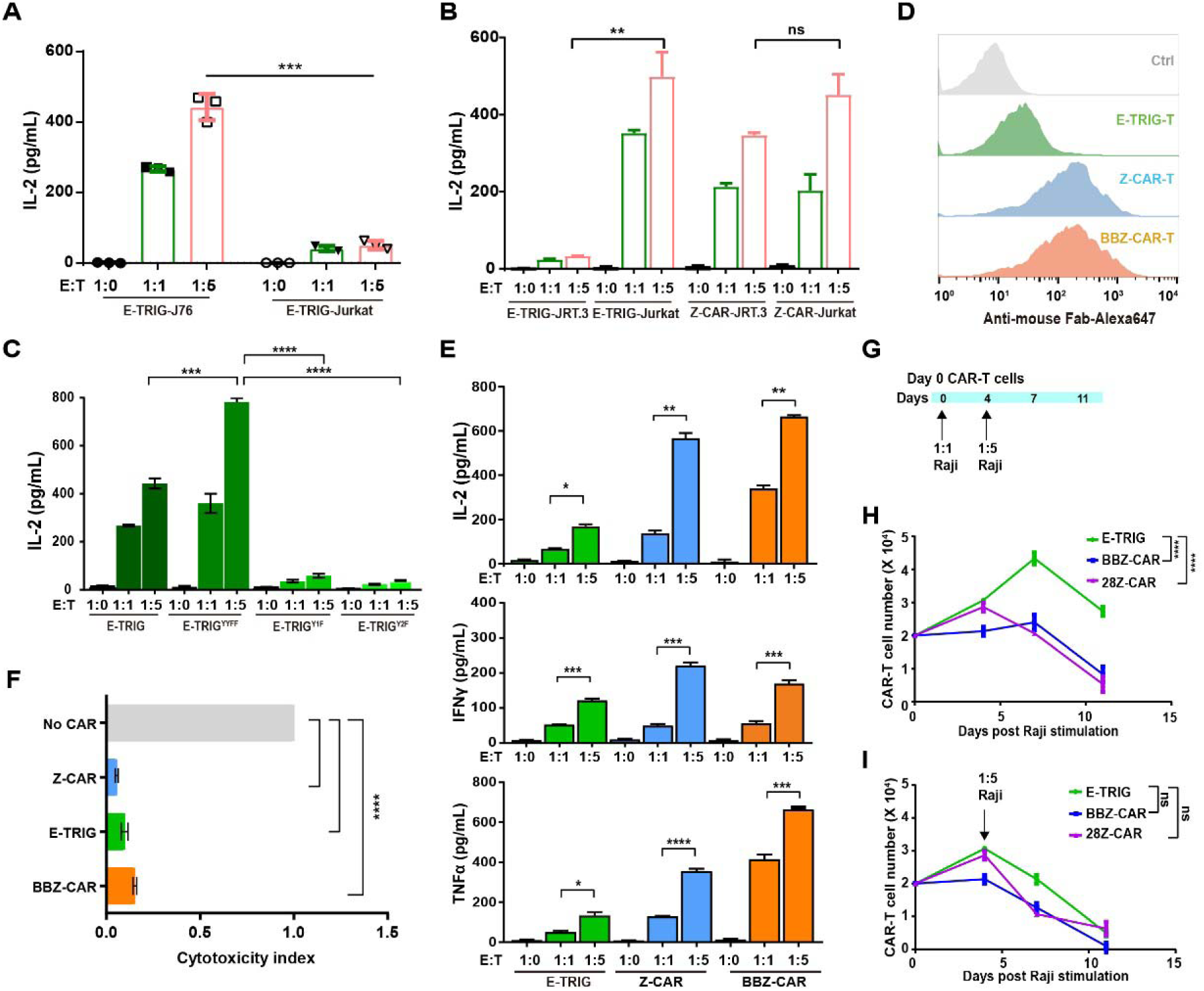
Endogenous TCR complex is indispensable for the activation of CD3ε based TRIG-T cells regardless of CD3ε ITAM motif. **A.** IL-2 secretion of E-TRIG engineered Jurkat or J76 cells cocultured with Raji cells at different E:T ratios detected by ELISA. Data were shown as Mean±SD (n=3; Unpaired t test; ns, *P* > 0.05; **, *P* < 0.01); **B.** IL-2 secretion of E-TRIG or Z-CAR engineered Jurkat or JRT3 cells cocultured with Raji B cells at different E:T ratios (n=3; Unpaired t test; ns, *P* > 0.05; \*\**P* < 0.01); **C.** IL-2 secretion of E-TRIG, E-TRIG^YYFF^, TRIG^Y1F^ or TRIG^Y2F^ engineered Jurkat cells cocultured with Raji cells at different E:T ratios (n=3; Unpaired t test; ***, *P* < 0.001; ****, *P* < 0.0001); **D.** Membrane surface expression of E-TRIG-T, Z-CAR-T and BBZ-CAR-T detected by flow cytometry; **E.** Various cytokines expression of E-TRIG, Z-CAR, or BBZ-CAR engineered primary T cells cocultured with Raji B cells at different E:T ratios (n=3; Unpaired t test; *, *P* < 0.05; **, *P* < 0.01; ***, *P* < 0.001; ****, *P* < 0.0001); **F.** The tumor cell-killing capacity of different CAR-Ts. E-TRIG, Z-CAR, or BBZ-CAR engineered primary T cells were cocultured with Raji-mCherry cells (n=3; Unpaired t test; ****, *P* < 0.0001); **G.** Experimental design showing cocultures with CAR-Ts and Raji cells at E:T ratio of 1:1 on day 0 followed by 1:5 E:T ratio on day 3 (n=3); **H and I.** CAR-T cells expansion in all groups were quantified by counting CAR-T cell numbers by fluorescent microscope (n=3NDs) (Two-way ANOVA with Turkey’s multiple-comparisons test was used to assess significance; ns, *P* > 0.05; ****, *P* < 0.0001).

Next, we measured the CARs function in human primary T cells. Pan CD3^+^ T cells were purified from the peripheral blood mononuclear cells (PBMC) and infected with lentiviruses of CARs (**Figure 3D**). These CARs were cocultured with Raji B cells and the secretion of cytokines were detected. Compared with the CD3Z-CAR or BBZ-CAR, CD3Ε-TRIG-T exhibited a milder functional profile, as evidenced by weaker but detectable secretion of IL-2, IFN-γ, and TNF-α (**Figure 3E**). Cytotoxicity of CAR-T cells were tested through coculture with Raji at an E:T ratio of 5:1 (**Figure 3F**). To mimic the clinical relapsed/refractory (R/R) chronic lymphocytic leukemia patients, we performed long-term coculture with Raji B cell and rechallenge the CAR-T cells with Raji B cell at a high E:T ratio (**Figure 3G**). CD3Ε-TRIG outperformed CD3Z-CAR and BBZ-CAR under these conditions (**Figure 3H and I**). Together, CD3Ε-TRIG-T produced less cytokines than CD3Z-CAR-T and BBZ-CAR-T, yet showed higher proliferation efficiency during long-term chronic antigen stimulation.

### CD3ε-TRIG-T cells have comparable therapeutic functions to BBZCAR-T cells in treating liquid tumors and CD3ε-TRIG-T could lead to a proliferative, memory-retaining, and exhaustion-reduced state in vivo

To investigate the CD3ε-TRIG function in vivo, we established the liquid tumor in NSIG mice using Nalm-6 cell (**Figure 4A**). In a standard mouse leukemia model, where 5 x 10^5^ CD19-expressing Nalm-6-luciferase-p2A-mCherry cells were injected, CD3ε-TRIG-T cells showed comparable therapeutic functions to BBZ-CAR-T cells compared with the PBS treated group (**Figure 4B and 4C**). Mice treated with CARs showed improved survival rate than PBS group (**Figure 4D**). Mice peripheral blood were collected at day 7 and day14, respectively. CD3ε-TRIG-T cells had greater expansion compared with the BBZ-CAR-T cells (**Figure 4E and Figure S4A**). BBZ-CAR-T cells upregulated the exhaustion markers PD-1, TIM-3, and LAG-3 than CD3ε-TRIG-T cells at day 14 (**Figure 4F and Figure S4B**). In addition, CD3ε-TRIG-T cells showed higher expression of CD45RO^+^ and CD62L^+^ central memory cells at day 21 (**Figure 4G and Figure S4C**). Meanwhile, tumors cells were reduced in blood treating with both CAR-T cells (**Figure 4H and Figure S4D**). In summary, these data suggest that CD3ε-TRIG-T cells exhibit a more persistent, central memory-prone, and exhaustion marker-reduced state.

**Figure 4.**
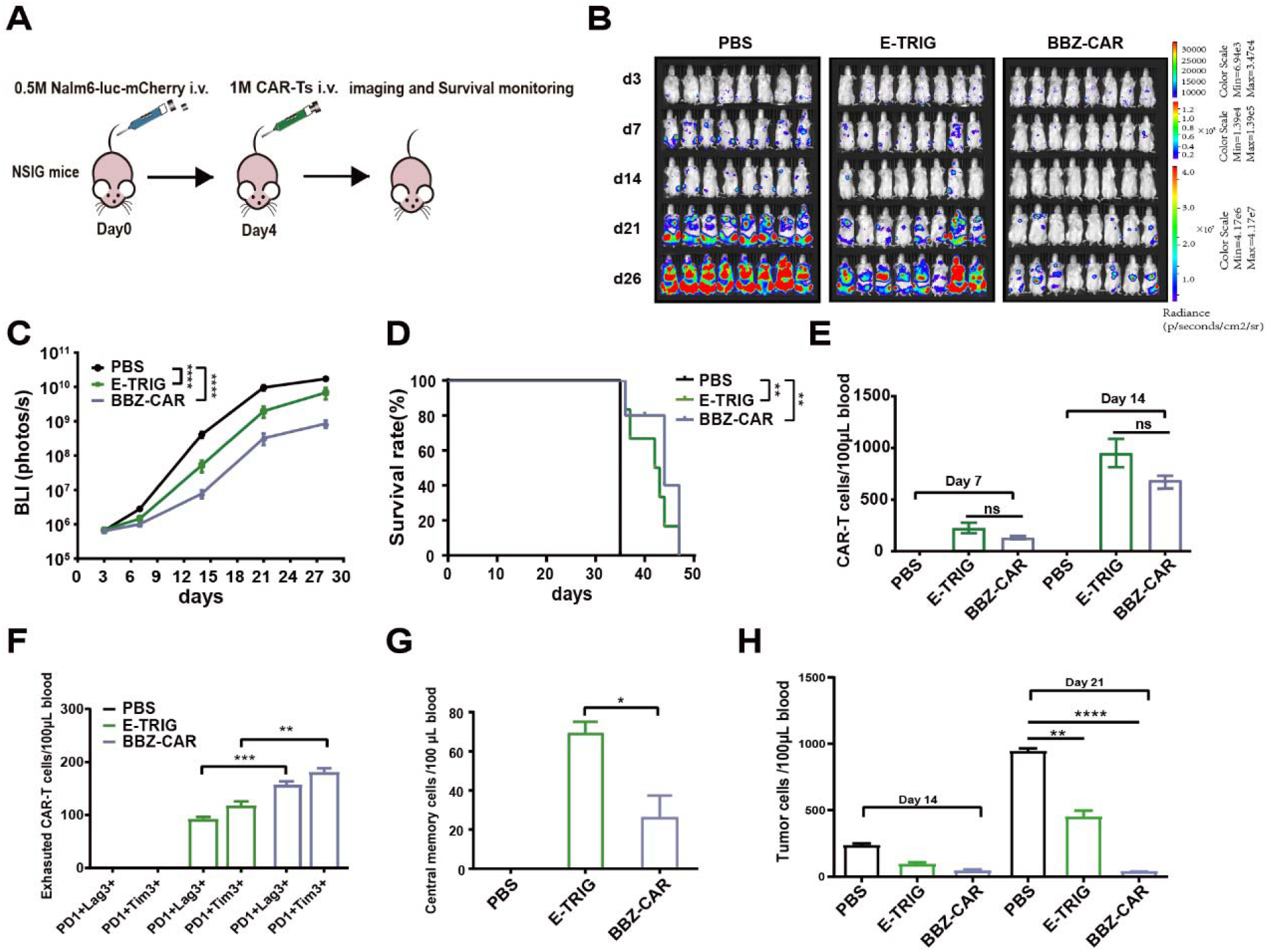
TRIG-T cells exhibit a proliferative, memory-retaining, and exhaustion-reduced phenotype in vivo. **A.** Schematics of the in vivo experimental workflow; **B.** Representative bioluminescence imaging of mouse model after CAR-Ts treatment. For all group, n=8 mice; **C.** Quantification of **B** showing the tumor burden as indicated by total radiance (p s^-1^ cm^-2^ sr^-1^). Two-way ANOVA with Turkey’s multiple-comparisons test was used to assess significance (****, *P* < 0.0001); **D.** Survival curve of NSIG mice in **B.** Mantel-Cox test was used to assess significance (**, *P* < 0.01); **E.** CAR-T cells expansion in vivo at day7 and 14 after injection determined via flow cytometry (Unpaired t test; ns, *P* > 0.05); **F.** Flow cytometry analysis of exhaustion marker PD1, TIM3 and LAG3 on CAR-T cells at day14 after injection (Unpaired t test; **, *P* < 0.01; ***, *P* < 0.001); **G.** Central memory markers expression on CAR-T cells at day 21 after injection (Unpaired t test; *, *P* < 0.05); **H.** Flow cytometry analysis of tumor cells in blood (Unpaired t test; **, *P* < 0.01; ****, *P* < 0.0001).

### In-silico design of TRIG molecules for T cell engineering

Inspired by the trigger mechanism of TCR, we designed a series of immuno-triggering peptides capable of phase separation with Lck and recruitment of the CD3ε intracellular domain (**Figure S5A**). These peptides exhibit phase separation ability with both Lck and CD3ε and can be categorized into two classes: those that strongly aggregate with Lck (T1, T3, T5) and those that exhibit strong self-aggregation (T2, T4, T6). Protein aggregation assays on two-dimensional membranes and protein condensation assays in three-dimensional solution showed that T5 possesses the strongest phase separation ability with Lck on 2D membranes (**Figure 5A and B**), whereas T6 forms the most robust self-aggregates under 3D conditions (**Figure S5B and C**). Moreover, both T5 and T6 efficiently recruited CD3ε on the two-dimensional membrane (**Figure 5C**). Using these peptides as the intracellular domains of immuno-triggering receptors, connected via the CD8α transmembrane domain to an extracellular anti-CD19 single-chain antibody, we designed a series of immuno-triggering receptors. Flow cytometry confirmed that all these receptors were well expressed on the T cell membrane (**Figure S5D**). IL-2 secretion results demonstrated that T cells engineered with these immuno-triggering receptors were significantly and specifically activated by CD19-expressing Raji cells (**Figure 5D**).

**Figure 5.**
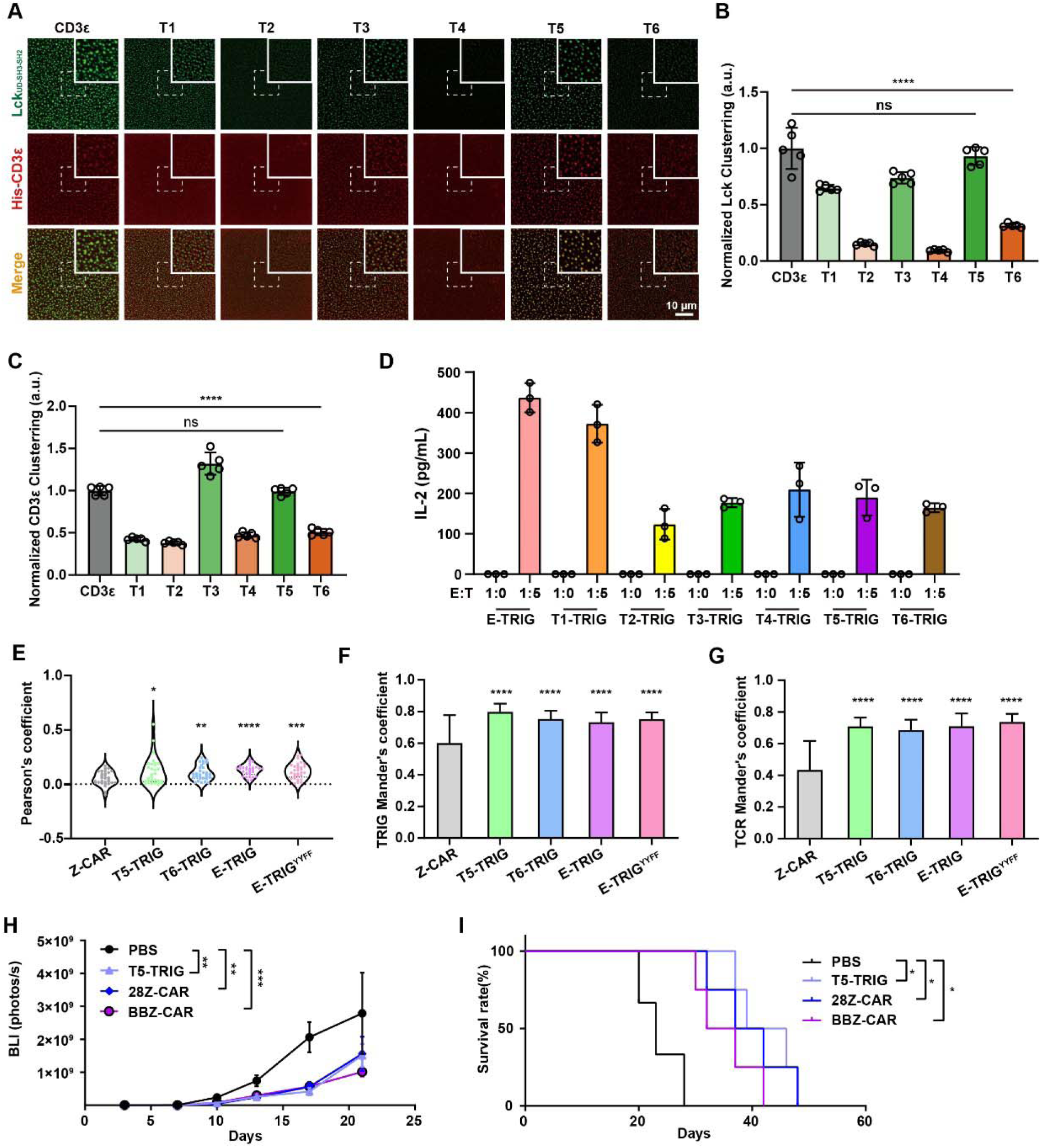
In silico design of TRIG molecules for T cell engineering. **A.** Typical images of the phase separation of Lck_UD-SH3-SH2_ (labelled with AF-488) and membrane-bound CD3ε (labelled with AF-647) on SLB. Phase separation was triggered with soluble CD3ε and T1-T6, Scale bar: 10μm; **B and C**. Normalized Lck (**B**) and CD3ε (**C**) clustering on SLB, which were calculated as described in Materials and Methods (Unpaired t test; ns, *P* > 0.05; ****, *P* < 0.0001); **D**. IL-2 secretion of TRIGs-engineered primary T cells stimulated with Raji cells, Data were shown as Mean ±SD, n=3; **E**. Pearson’s coefficient in colocalized volume in Figure S5E. Significance comparisons are shown for all groups against the Z-CAR group (n=30; Unpaired t test; *, *P* < 0.05; **, *P* < 0.01; ***, *P* < 0.001; ****, *P* < 0.0001); **F-G**. Histogram of Manders’ coefficient of TRIG (**F**) and TCR (**G**) molecules in Figure S5E. Significance comparisons are shown for all groups against the Z-CAR group, (n=30; Unpaired t test; ****, *P* < 0.0001); **H**. Quantification of PBS, T5-TRIG, 28Z-CAR and BBZ-CAR groups in Figure S5G showing the tumor burden as indicated by total radiance (p/s). Two-way ANOVA with Tukey’s multiple-comparisons test was used to assess significance (n=6; **, *P* < 0.01; ***, *P* < 0.001); **I**. Survival curve of PBS, T5-TRIG, 28Z-CAR and BBZ-CAR groups in Figure S5G. Mantel–Cox test was used to assess significance (n=6; *, *P* < 0.05).

Given that these receptor intracellular domains lack the ability to directly transduce downstream signals, we hypothesize that they activate T cell immune functions by recruiting endogenous TCR molecules, thereby initiating T cell activation. To test this hypothesis, we immobilized CD19 on an SLB surface to stimulate Jurkat cells expressing chimeric antigen receptors (CD3ε, CD3ε-YYFF, T5, T6, CD3ζ) and analyzed the distribution of the trigger and TCR by TIRF-SIM imaging. Upon SLB-CD19 stimulation, the trigger formed clusters and, to some extent, recruited endogenous TCR enriched around the clusters (**Figure S5E**). Quantitative analysis using both Pearson’s correlation coefficient and Manders’ coefficient between the trigger and endogenous TCR revealed that the trigger significantly recruited endogenous TCR, whereas the CD3ζ-CAR showed little ability to do so (**Figure 5E and G**). Moreover, when using CD19 immobilized on a glass-bottom surface for stimulation, we observed that TRIGs mediated stronger self-clustering compared to CD3ζ-CAR in Jurkat cell (**Figure S5F**). Cellular functional experiments and imaging data both indicated that TRIGs might confer superior anti-tumor efficacy. Furthermore, we evaluated their performance in mouse tumor model. We injected Raji-luciferase-p2A-mCherry cells into NSIG mice to establish a humanized hematological tumor model and treated the mice with various CAR-T cells. T5-TRIG exhibited tumor clearance comparable to that of BBZ-CAR or 28Z-CAR and resulted in better mouse survival (**Figure 5H and I, Figure S5G and I**).

### Harnessing phase separation-based trigger mechanism endows NK cell liquid tumor killing ability

NK cells and T cells are closely related in terms of cell differentiation and share some similarities in immune functions (Tognarelli et al., 2023). NK cells can induce an antigen-independent immune response against malignant cells and many studies have shown promising anti-tumor effects when using NK cell-based immunotherapy (Fang et al., 2017). Moreover, several activating receptors on NK cell membranes possess complex structures analogous to the T-cell receptor (TCR), including NCR1, NCR3, CD16, KIR3DS1, and NKG2D. These receptors consist of a single-pass transmembrane receptor and associated adaptor proteins (such as CD3ζ, FcRγ, DAP10, and DAP12) that couple via transmembrane domains. We hypothesize that within these receptor complexes, some receptors may also utilize a mechanism involving phase separation between the receptor intracellular domain and kinases to trigger transmembrane signal transduction.

We analyzed several activating single-pass transmembrane receptors on NK cell membranes, including NKG2D, KIR3DS1, NCR1, NCR3, and CD16α. Two-dimensional membrane protein clustering assays revealed that these receptors exhibit varying levels of phase separation with Lck (**Figure 6A and B**). Like CD3ε-TRIG, we designed novel TRIG molecules incorporating the intracellular domains of these NK activating receptors. Flow cytometry analysis of chimeric NK receptor expression on the membrane of NK92MI cells showed that all receptors, except those containing the CD3ε and NCR3 intracellular domains, displayed high surface expression levels (**Figure S6A**). When NK92MI cells transduced with these chimeric NK-triggering receptors were co-cultured with CD19-expressing Nalm6 cells, specific activation-induced cytokine secretion was detected. Notably, the CD3ε intracellular domain elicited the highest activation level, while the NKG2D and KIR3DS1 intracellular domains, which exhibit strong phase-separation ability, also induced robust NK cell immune activation (**Figure 6C and D**). The TRIG constructs (T1-T6), previously validated in T cells, were also transduced into NK92MI cells, and their membrane expression levels were assessed by flow cytometry (**Figure S6B**). The ability of these TRIGs to activate NK cells was measured by cytokine secretion in co-culture assays (**Figure 6E and F**). Among them, T5 demonstrated excellent performance in both membrane expression and NK cell activation. Next, we evaluated the therapeutic efficacy of immune-triggering receptor-engineered NK cells in a mouse tumor model (**Figure 6G**). Mice bearing Raji-luciferase-p2A-mCherry cells were treated with various engineered NK cells. We observed that TRIG-NK cells significantly eliminated tumor cells and suppressed tumor growth in vivo (**Figure 6H, Figure S6C**). Analysis of peripheral blood NK cell proportions in surviving mice on day 35 revealed a higher percentage of NK cells in T5-NK-treated mice, suggesting that T5 promoted more sustained NK cell survival and proliferation (**Figure 6I**). TRIG-NK treatment substantially prolonged the survival of tumor-bearing mice (**Figure 6J**).

**Figure 6.**
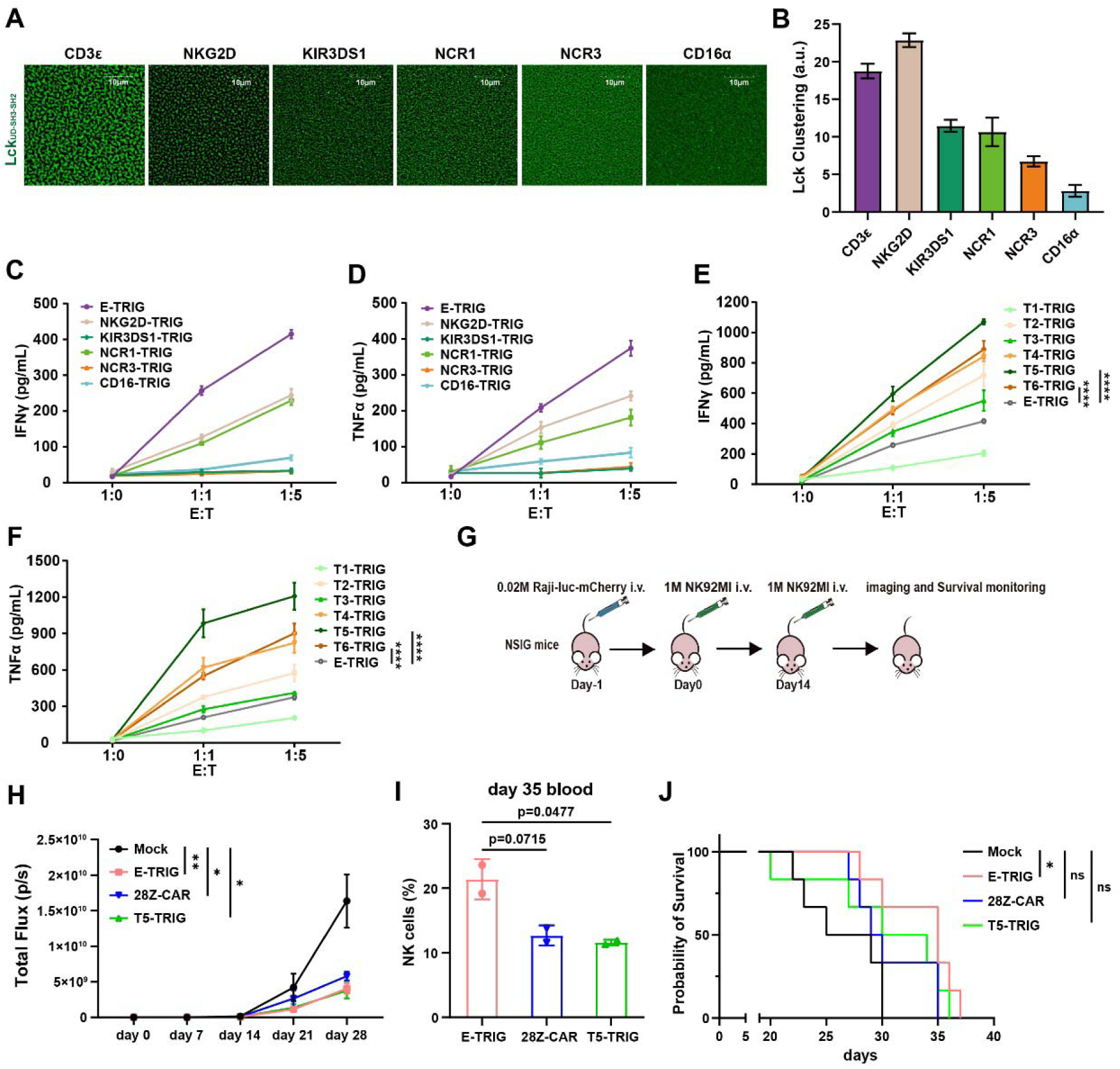
Harnessing phase separation-based trigger mechanism endows NK cell liquid tumor killing ability. **A** and **B**. Typical images (**A**) and analysis (**B**) of NK receptors induced clustering of Lck, Scale bar: 10μm; **C** and **D**. IFN-γ (**C**) and TNF-α (**D**) secretion of NK receptors-TRIGs engineered NK92MI cells stimulated with Nalm6 cells, Data were shown as Mean±SD; **E** and **F**. IFN-γ (**E**) and TNF-α (**F**) secretion of T1-T6 and E-TRIGs engineered NK92MI cells stimulated with Nalm6 cells, Data were shown as Mean±SD (n=3; Unpaired t test; ****, *P* < 0.0001); **G.** Schematics of the in vivo experimental workflow of TRIG-NK cells; **H.** Quantification of Figure S6C showing the tumor burden as indicated by total radiance (p/s). Two-way ANOVA with Tukey’s multiple-comparisons test was used to assess significance (n=6; *, *P* < 0.05; **, *P* < 0.01); **I.** Quantification of TRIG-NK cell proportions in mouse peripheral blood lymphocytes on day 35, n=2; **J.** Survival curve of NSIG mice in Figure S6C. Mantel–Cox test was used to assess significance (n=6; ns, *P* > 0.05; *, *P* < 0.05).

Furthermore, we aim to verify the recruitment of endogenous activating receptors by TRIG via phase separation during TRIG-mediated NK cell activation. Flow-cytometry with fluorescent antibody labeling was used to assess the surface expression levels of NCR1, NCR3, and NKG2D on NK92MI cells (**Figure S6D**). To verify that TRIGs recruit endogenous NK activating receptors, we immobilized CD19 on supported lipid bilayers (SLBs) to stimulate NK92MI cells expressing chimeric triggering receptors (T5, T6, or CD3ζ), and analyzed the clustering of endogenous NK receptors using TIRF-SIM imaging. Under stimulation with CD19 immobilized on SLBs, the TRIG constructs induced greater clustering of endogenous NK receptors NCR1, NCR3, and NKG2D compared to CD3ζ (**Figure S6E–G**). These findings suggest that TRIG initiates NK cell immune responses by recruiting endogenous activating receptors. Additionally, we found that different TRIGs exhibit preferential recruitment toward distinct endogenous receptors in NK cells.

## Discussion

Discoveries in TCR signaling have the potential to profoundly influence the design of CARs and TCR-like chimeric receptors, advancing their effectiveness in cancer immunotherapy. The design of chimeric antigen receptors is mostly based on the intracellular domain of the CD3ζ subunit in the TCR receptor complex as an important component for signal transduction of the CAR molecule, for downstream signal activation and related protein expression, as well as for the execution of killing of target cells. However, this design makes the signal richness of CAR much lower than that of a true TCR. On one hand, this may lead to unregulated CAR signaling, where overly strong early signals trigger cytokine storms and neurotoxicity; on the other hand, it may result in a poorer memory phenotype of CAR-T cells, preventing long-term efficacy (Yang et al., 2026). Increasing the diversity of the intracellular domains of CAR molecules can improve the long-term therapeutic effect of CAR-T cells to some extent. Additionally, approaches like TRuC-T and STAR-T redesign the extracellular domains of TCRs to directly utilize all TCR signaling components, allowing the modified T cells to maintain both signal richness and antigen recognition diversity.

To better harness the full TCR signaling machinery in the engineering design of T cells, a deeper understanding of the TCR activation mechanism is required. TCR engagement with pMHC leads to phosphorylation of its intracellular domains by kinases, thereby transmitting signals downstream. Extensive structural biology studies have shown that the extracellular domains of the TCR undergo no marked conformational changes upon pMHC binding; therefore, the conventional allosteric model may not adequately explain TCR activation. Considering the mechanical force imposed by pMHC binding, mechanical force-induced conformational changes in the TCR are gaining increasing attention in the field. Although the structure of the CD8/TCR/pMHC ternary complex remains unresolved, studies have demonstrated that pMHC binding is enough to activate T cells even in the absence of CD8 signaling, in which process free Lck may play a critical role (Wei et al., 2020). Conformational changes in the intracellular domains upon TCR activation are now widely accepted. Our findings, together with previous studies, have shown that the intracellular domains of the CD3ε and CD3ζ subunits are sequestered in a membrane-proximal state in resting T cells and are released from the membrane upon ligand engagement. Here, we further found that the intracellular domains of CD3δ and CD3γ remain constitutively away from the plasma membrane. These results indicate that the release of the CD3ε and CD3ζ intracellular domains from the membrane following pMHC binding represents a key initiating event in TCR triggering. In addition, although both CD3ε and CD3ζ can associate with the membrane in the resting state, their binding dynamics differ: CD3ζ is maintained at a low level of phosphorylation in resting T cells, suggesting that its signaling specificity is relatively low, which may be related to the requirement for tonic TCR signals to sustain T cell survival. Collectively, these findings suggest that CD3ε responds with greater specificity to ligand engagement.

The strength and duration of TCR-pMHC interaction also critically influence TCR activation. Low-affinity pMHC binding may fail to induce enough release of the CD3ε intracellular domain from the plasma membrane, while a short-lived binding event may prevent the stable maintenance of the membrane-distal conformation of CD3ε, both of which suppress CD3ε phosphorylation. Moreover, under pathological conditions, activating pMHC molecules are often present at very low abundance; tumor cells, for instance, may express only a few to a few dozen such complexes. T cells use TCRs enriched at the tips of microvilli to rapidly scan target cells, facilitating the search for rare activating pMHC ligands (Cai et al., 2017). However, TCR phosphorylation triggered by a limited number of pMHC engagements is also constrained by abundant negative regulators on the plasma membrane, such as CD45. In our previous work, it was found that the CD3ε chain can interact with Lck through ionic interactions, leading to Lck/TCR condensation, thereby upregulating Lck activity and enhancing Lck-mediated CD3ε phosphorylation. The CD3ε/Lck clusters can recruit more TCR and Lck to promote TCR signaling, and then forming a self-promoting system (Chen et al., 2023). The intracellular condensation cooperates with extracellular size exclusion facilitate TCR to overcome the inhibition and generate enough signal to initiate LAT signaling (Chen et al., 2023). In this study, we found CD3ε-incorparated CAR can mediate T cell activation, mechanical force but not phase separation mediated clustering are essential in triggering CD3ε activation. Mechanical force loading on TCR may modulate the conformation of TCR transmembrane domain, and enhancing the local membrane tension, resulting the release of CD3ε-ICD from plasma membrane. Collectively, these findings suggest that CD3ε is the principal mechano-responsive element within the TCR, which amplifies proximal TCR signaling through force-induced phase separation to reach the activation threshold and trigger downstream LAT signaling. Thus, beyond its signal-modulating function, CD3ε also serves as a critical signaling “switch” within the TCR complex.

Many immune cells express receptor complexes assembled from multiple single-pass transmembrane proteins, analogous to the TCR, such as the B cell receptor (BCR) and various receptors on NK cells (Chen et al., 2024). Several NK cell receptors not only share a similar assembly mode with the TCR but also contain the CD3ζ subunit (Sivori et al., 2019). In these receptors, CD3ζ is considered the principal signaling module, whereas the intracellular domains of the cognate receptor chains generally lack canonical signaling motifs, such as ITAMs, and their specific functions have remained unclear (Sivori et al., 2019). We found that the intracellular domains of these receptors can also undergo phase separation with Lck, and that, like CD3ε, they can recruit endogenous receptors to trigger NK cell activation. We therefore hypothesized that triggering receptor-Lck phase separation represents a potential strategy for harnessing endogenous receptors to activate immune cells, and we term this approach TRIG. We designed a series of TRIG molecules that can phase-separate with Lck but lack intrinsic signaling motifs, and found that these TRIG molecules can recruit the corresponding endogenous receptors to mediate the activation of both T cells and NK cells. In relevant mouse tumor models, they exhibited therapeutic efficacy comparable to that of second-generation CARs. These results suggest that designing intracellular domains with the ability to undergo phase separation with a kinase can achieve selective receptor activation and thereby modulate immune cell functions, offering a potential new strategy for immune cell engineering.

Traditional CAR designs predominantly rely on CD3ζ for signal transduction, with additional modulation by various costimulatory domains, which to some extent restricts the diversity of CAR-T signaling. Recent studies have directly compared the functions of the four CD3 intracellular domains using second-generation CAR platforms based on CD28 and 4-1BB, revealing that CAR T cells harboring the CD3ε intracellular domain outperform CD3ζ CAR T cells in vivo (Velasco Cárdenas et al., 2023; Wang et al., 2025). CD3ε has also been directly incorporated into conventional 28Z-or BBZ-CARs, where it was found can regulate TCR signaling and T cell function by recruiting Csk (Wu et al., 2020; Xu et al., 2024). Furthermore, recent studies show that CD3δ and CD3ζ can each recruit the inhibitory phosphatase SHP-1 to modulate TCR signal strength (Gaud et al., 2023; Velasco Cárdenas et al., 2023). CD3ζ contains three ITAMs whose phosphorylation occurs in a hierarchical manner, with distinct phosphorylation patterns giving rise to different downstream signals (Feucht et al., 2019; Li et al., 2025). Thus, for an intact TCR, both the types of ITAMs and their phosphorylation patterns are highly diverse, enabling the TCR not only to mediate T cell activation but also to govern T cell fate through a “phosphorylation code” (Chen et al., 2025). Consequently, relying solely on CAR-intrinsic signaling cannot achieve the same level of fine regulation as the TCR. The TRIG molecules we designed here undergo phase separation with Lck and thereby recruit endogenous TCRs. This strategy achieves phosphorylation diversity on the one hand, and on the other, because TRIG molecules lack phosphorylation motifs, they do not antagonize endogenous TCR signaling. Since different TRIG molecules exhibit distinct phase separation properties with Lck, the phosphorylation patterns elicited by the recruitment of endogenous TCRs may also differ. The molecular mechanism of such “phase separation-driven phosphorylation” requires more in-depth investigation to enable precise encoding of T cell activation and differentiation through TRIG molecules.

Phase separation plays a critical role in T cell activation. T cells exploit phase separation to achieve proximal signal amplification from a limited number of activated receptors, such as the TCR and LAT, thereby enabling productive T cell activation. TCR condensation with Lck also promotes immunological synapse assembly, which enhances T cell killing capacity (Chen et al., 2023). This mechanism has been harnessed in CAR-T design: studies have shown that appending CD3ε to a second-generation CAR enhances CAR molecular clustering and promotes immunological synapse formation (Xu et al., 2024). Other studies have incorporated additional phase-separation-prone protein modules into second-generation CARs, which markedly augments CAR clustering and boosts CAR-T cytotoxic function (Zhang et al., 2026). However, the contribution of endogenous TCR signaling in these clustering-enhanced CAR-T cells remains poorly characterized, and whether these clustered CAR molecules can recruit other endogenous receptors is still unknown. Beyond directly altering intracellular domain distribution and signal transduction by adding phase separation elements to the CAR intracellular domain, an alternative approach has been to engineer the extracellular domain to achieve phase separation-mediated receptor activation (Li et al., 2022). Here, we found that inducing phase separation at the extracellular domain of the CAR molecule fails to directly activate CAR-T cells. This is possibly due to the relatively long linker region of the CAR and the strong membrane association of CD3ε, which may prevent the propagation of extracellular clustering into conformational changes at the intracellular membrane-proximal state, a mechanism fundamentally distinct from that of the TCR complex. Nevertheless, incorporating environmentally sensitive phase separation modules into the CAR extracellular domain enables conditional clustering of CAR molecules, which, in the presence of immobilized ligand binding, can enhance CAR-T cell function under specific microenvironmental conditions.

While CAR-T cells have demonstrated remarkable success in hematologic cancers by recognizing surface antigens independent of presentation, their efficacy in solid tumors remains limited due to challenges such as immunosuppressive microenvironments, poor trafficking, and antigen heterogeneity (Zugasti et al., 2025). In contrast, TCR-based therapies leverage the natural precision of MHC-restricted antigen recognition, offering potential against intracellular targets but facing hurdles related to MHC downregulation and off-target toxicity (Chen and Zhou, 2026). In this study, we identified a trigger mechanism of TCR activation in T cell, and designed new TRIG molecules to mediate the activation of T cell and NK cell which depends on endogenous receptor signaling. Despite their current functional limitations, TRIG molecules exploit an endogenous receptor–dependent mechanism of immune cell activation, giving them substantial research value for future therapies against solid tumors and other diseases and warranting further investigation and optimization.

## Data Availability Statement

All data generated or analyzed in this study are included in the article and the Supplementary Information document.

## Acknowledgement

We acknowledge Prof. Pilong Li (Tsinghua University) for kindly providing the related plasmid for the phase separation system (SIM6-PV and SUMO5). We thank Prof. Yang Yu for providing the firefly-luciferase expression plasmid. We thank Prof. Pengcheng Bu’ lab (Institute of Biophysics, Chinese Academy of Sciences) for technical supporting of the animal assay. We thank Dr. Yun Feng and Qing Bian for the help in TIRF-SIM (Total Internal Reflection Fluorescence-Structured Illumination Microscopy) imaging and data processing with Imaris. All the imaging experiments were performed at the Center for Biological Imaging, Institute of Biophysics, Chinese Academy of Science. We thank Dr. Junying Jia and Dr. Shu Meng (Core Facility, Institute of Biophysics, CAS) for technical support in the flow cytometry analysis. We thank Yihui Xu (State Key Laboratory of Biomacromolecules, Institute of Biophysics, Chinese Academy of Sciences) for in vivo imaging technology.

## Author contribution

S.W., H.C., and J.L. designed the experiments. H.C., C.L., W.C. and J.L. conceived the project. S.W. constructed the cell lines, performed the cell function experiments and in-vivo assays. H.C. performed phase separation studies and FRET experiments. Q.W. performed cell function experiments and CAR-NK assays. W.H. participated in protein purification. S.W., H. C., and Q.W. prepared initial figures and manuscript. X.G., and H.Q. participated in the plasmid construction and animal assays. All authors helped revise the manuscript.

## Materials and Methods

### Antibodies

Antibodies used in this study were purchased from various commercial suppliers: Biotinylated anti-human CD3ε (UCHT1, Abcam, ab191112); Anti-Human CD279(PD1)-SB436 (Invitrogen, 62-2799-42); Anti-human CD69-APC (SinoBiological, 11150-MM06-A); Anti-human CD45-APC (Biolegend, HI30); Anti-human CD366(TIM3)-APC (Invitrogen, 17-3109-42); Anti-Human CD233(LAG-3)-PE-cyanine7(Invitrogen,25-2239-42); Anti-Human CD45RO-SB436 (Invitrogen, 62-0457-42); Anti-Human CD62L-BV650(Invitrogen, 416-0629-42); Anti-Human CD3ε-PE (Invitrogen, 12-0038-42); anti-β2M (Abcam, AB187602); Biotinylated anti-huamn CD28 (CD28.2, Abcam, ab267385); anti-mouse Fab2-Alexa647 (Jackson Immunoresearch, 115-606-072); Anti-Human NCR1 (Invitrogen, PA5-102860); Anti-Human NCR3 (Invitrogen, PA5-104376); Anti-Human NKG2D (Invitrogen, PA5-102038).

### Cell lines and lentiviral transfection

All lentivirus constructs were generated by inserting target coding sequences into lentivirus transfer plasmid backbone (pHAGE-P2A-zsgreen or pHAGE-P2A-mCherry). Lentiviral constructs were co-transfected into HEK293T cells with helper plasmids pMD2.G and psPAX2 using Polyplus transfection reagent (Polyplus, #101000046). Viral supernatant was harvested at 48 and 72 h of transfection.

CAR-Jurkat/NK92MI cells were generated by infecting Jurkat/NK92MI cells with lentivirus encoding CAR constructs, followed by quantification of surface CAR expression with anti-Mouse Fab-Alexa647 staining and flow cytometry analysis. JRT3 cell line was purchased from ATCC. Human cancer cell line Nalm6 was purchased from Procell (Wuhan). J76 and NK92MI cell line was kindly provided by Dr. Wei Chen (Zhejiang University). All Jurkat cell lines (Jurkat, J76, and JRT3) and B cells, Raji and Nalm6, were cultured in RPMI-1640 medium supplemented with 100 U/mL streptomycin, 100 ug/mL penicillin and 10% fetal bovine serum at 37℃ with 5% CO_2_. NK92MI cells were cultured in MEM supplemented with 100 U/ml IL-2, 12.5% FBS, 12.5% Horse serum, 100 ug/mL penicillin, 0.2 mM Myo-inositol, 0.02 mM Folic acid, and 0.1 mM β-mercaptoethanol. All cells were tested regularly for mycoplasma contamination and were negative.

### Peptides and proteins

#### Synthesized peptides

All peptides used in this study were synthesized by GL biotech (Shanghai). Fluorescence dye labeling was conducted as previously described (Chen et al., 2023).

#### Lck protein

The expression and purification of Lck proteins were conducted as previously described (Chen et al., 2023). Briefly, Lck_UD-SH3-SH2_ (residues 3-226) were expressed in BL21 (DE3) *E. Coli* (Tsingke) and then purified with His-Trap HP column followed by gel filtration (Hiload 16/600, Superdex 200, Cytia).

#### SIM6-PV and SUMO5

The genes encoding SIM6-PV and SUMO5 were kindly provided by Prof. Pilong Li (Tsinghua University). The genes were subcloned into a home-modified pCOLD vector with an N-terminal GST tag followed by a 3C protease cleavage site. The recombinant plasmids were transformed into BL21(DE3) *E. Coli* (Tsingke) cells. The cells were cultured at 37 ℃ until the optical density reached the mid-log phase, and protein expression was induced with 0.5 mM IPTG at 15 ℃ for 12 h. After cell lysis, the supernatant was collected and incubated with a glutathione affinity column at 4 ℃ for 2 h. The GST tag was subsequently removed by on-column cleavage using 3C protease. The proteins were then concentrated and further purified by gel filtration chromatography (HiLoad 16/600 Superdex 75, Cytiva).

### Protein condensation assays on supported lipid bilayer (SLB)

SLB formation and protein condensation assays were performed as previously described (Du et al., 2025). Briefly, small unilamellar vesicles (SUVs) were applied to pre-cleaned glass-bottom 96-well plates and incubated at 37 °C for 1 h to form lipid bilayers. His-tagged proteins were ligated to the pre-formed SLBs, and the passenger molecules were then added to trigger on-membrane phase separation. For the membrane tension modulation assay, 5 mM MβCD was injected to extract cholesterol from the pre-formed SLB, after which CD3ε or phosphorylated CD3ε was added to initiate phase separation. All imaging was performed using an Olympus FV1200 microscope equipped with a 60× oil-immersion objective.

Co-localization of CD3/Lck/TRIGs clustering on SLB were quantified with confocal imaging as previously reported (Chen et al., 2023). In brief, His-tagged CD3ε and Lck constructs were incubated with preformed SLB in sequence. Soluble TRIGs cytoplasmic tail peptides were added into the reaction chambers to initiate phase separation. Confocal microscopy was performed on an Olympus FV1200 system equipped with a 60× oil-immersion objective, protein clustering was quantified with Image J.

### GUV assay

GUVs were produced using a previously reported gel-assisted assay (Weinberger et al., 2013). Briefly, 50 μL of PVA solution (5% w/w) was added to a 96-well plate and dried for 1 h. Then, 15 μL of lipids (95% DOPC, 5% DGS-NTA-Ni^2+^) in chloroform (1 mg/mL) were spread onto the PVA film and placed under vacuum for 30 min. Next, 50 μL of PBS was added and incubated at room temperature for 1 h. The resulting GUVs were harvested and validated by confocal imaging. His-tagged CD3ε and His-tagged Lck_UD-SH3-SH2_ were ligated onto the GUVs, followed by osmotic adjustment (PBS for isotonic, ddH₂O for hypotonic, and sorbitol for hypertonic conditions). The GUV suspensions were then incubated at room temperature for 30 min and imaged using an Olympus FV1200 confocal microscope equipped with a 60× oil-immersion objective.

### Fluorescence resonance energy transfer (FRET) assay

FRET assays were performed as previously reported (Yang et al., 2017). Jurkat cells harboring mTFP-fused CD3-CARs were stained with R18 and then stimulated on glass surfaces coated with biotinylated proteins or antibodies (anti-CD3ε, anti-CD28, anti-mouse Fab). Biotinylated anti-human β2m was used as a control. mTFP was excited with a 450 nm laser and detected at 470–520 nm; R18 was excited with a 561 nm laser and detected at 570–650 nm. An image of a cell in contact with the surface was first acquired (before bleaching). The entire cellular R18 signal was then fully photobleached using the 561 nm laser at 100% power for 40–60 s, after which another image of the same cell was captured (after bleaching). FRET efficiency was calculated as (FL_AP_ – FL_BP_) / FL_AP_, where FL_AP_ represents the mTFP1 fluorescence after photobleaching and FL_BP_ represents the mTFP1 fluorescence before photobleaching. Imaging was performed on an Olympus FV3000RS confocal microscope.

### CAR-T cell preparation

Lentiviral particles contained CAR constructs were harvested and concentrated, and used to infect the target cells in the presence of 8 µg mL^−1^ Polybrene, followed by spinoculation at 1600 rpm for 60 min. CAR surface expression was validated using anti-Mouse Fab-Alexa647 (Jackson Immunoresearch, #115-606-072) and flow cytometry.

For primary CAR-T cells, pan CD3^+^ T cells were isolated from PBMCs (Milestone® Biotechnologies) using the MojoSort Human CD3 T cell isolation Kit (Biolegend, #480022). T cells were activated with Human T-activator CD3/CD28 Dynabeads (Thermo Fisher Scientific, #11161D) on day 0, transduced with CAR lentivirus at day 1, and expanded with media doubling and human IL-2 supplementation every 2-3 days. Dynabeads were removed on day 7 with magnetic separation. CAR-T cells were harvested and the transduction efficiency was assessed by ZsGreen and CAR surface staining.

### T/NK cell-Raji B co-culture

Generally, for cytokines secretion assays, Raji B cells were co-cultured with 2×10^4^ Jurkat cells in 1640 medium or with 2×10^4^ NK92MI cells in MEM medium for 18-24 hours. Supernatants was collected and analyzed using ELISA kit for cytokines quantification.

### Cytotoxicity assay

Cytotoxicity was measured by flow cytometry. Briefly, mCherry-expressing Raji B cells were cocultured with CARs engineered primary CD3^+^ T cells at an effector to target ratio of 5:1. After 24 hours of incubation, cells were collected and washed with PBS. Cells were then detected by flow.

Cytotoxicity index= live tumor cell percentage (CAR-T treated group) / live tumor cell percentage (Mock group)

### TIRF-SIM imaging

Co-localization of TRIGs with TCR or NK receptors in cells was quantified by TIRF-SIM imaging as previously described (Chen et al., 2023). Briefly, CD19 or SIM6-PV was coated onto pre-formed SLBs or glass surfaces. Cells stably expressing TRIGs were added and incubated at 37 °C for 40 min. The cells were then fixed, permeabilized, stained with fluorophore-conjugated antibodies, and imaged using a TIRF-SIM microscope. Image processing and data analysis were performed with ImageJ and Imaris 10 (Bitplane).

### Mouse Experiments

Female NSIG mice (6-8weeks, female, Beijing HFK Bioscience Corporation) were housed in the animal facility of the Institute of Biophysics, Chinese Academy of Sciences. Mice were injected intravenously with 0.5million Nalm6-firefly luciferase-p2A-mCherry cells, or Raji-firefly luciferase-p2A-mCherry cells, on day 0 and treated with 1 million CAR-T cells intravenously on day 4. Disease progression was monitored by bioluminescence imaging and survival analysis. For in vivo phenotyping studies, 2.5 µg human recombinant IL-2 was administered subcutaneously every 2 days. Blood was drawn on days 7, 14 and 21 after CAR-T injection for flow cytometry analysis of CAR-T percentage and the expression of exhaustion and central memory markers. CoutBright^TM^ Absolute counting Beads (Thermo Fisher) were used as an internal standard according to the manufacturer’s instructions to calculate absolute cells counts in cell suspensions. Disease progression was monitored by bioluminescence imaging and survival analysis. All procedures were approved by the Biomedical Research Ethics Committee of the Institute of Biophysics, Chinese Academy of Sciences (SYXK2023189), and conducted according to the institutional guidelines.

For NK cells therapy, mice were injected intravenously with 20 thousand Raji-firefly luciferase-p2A-mCherry cells on day -1 and treated with 1 million CAR-NK cells intravenously on day 1 and 14. Disease progression was monitored by bioluminescence imaging and survival analysis. 0.5 µg human recombinant IL-15 was administered subcutaneously twice a week. Blood was drawn on days 35 for flow cytometry analysis of CAR-NK percentage. CoutBright^TM^ Absolute counting Beads (Thermo Fisher) were used as an internal standard according to the manufacturer’s instructions to calculate absolute cells counts in cell suspensions. Disease progression was monitored by bioluminescence imaging and survival analysis. All procedures were approved by the Biomedical Research Ethics Committee of the Institute of Biophysics, Chinese Academy of Sciences (SYXK2023189), and conducted according to the institutional guidelines.

