## Supporting materials for "Intrinsic immune receptor triggering mechanism informs rational chimeric receptor design"

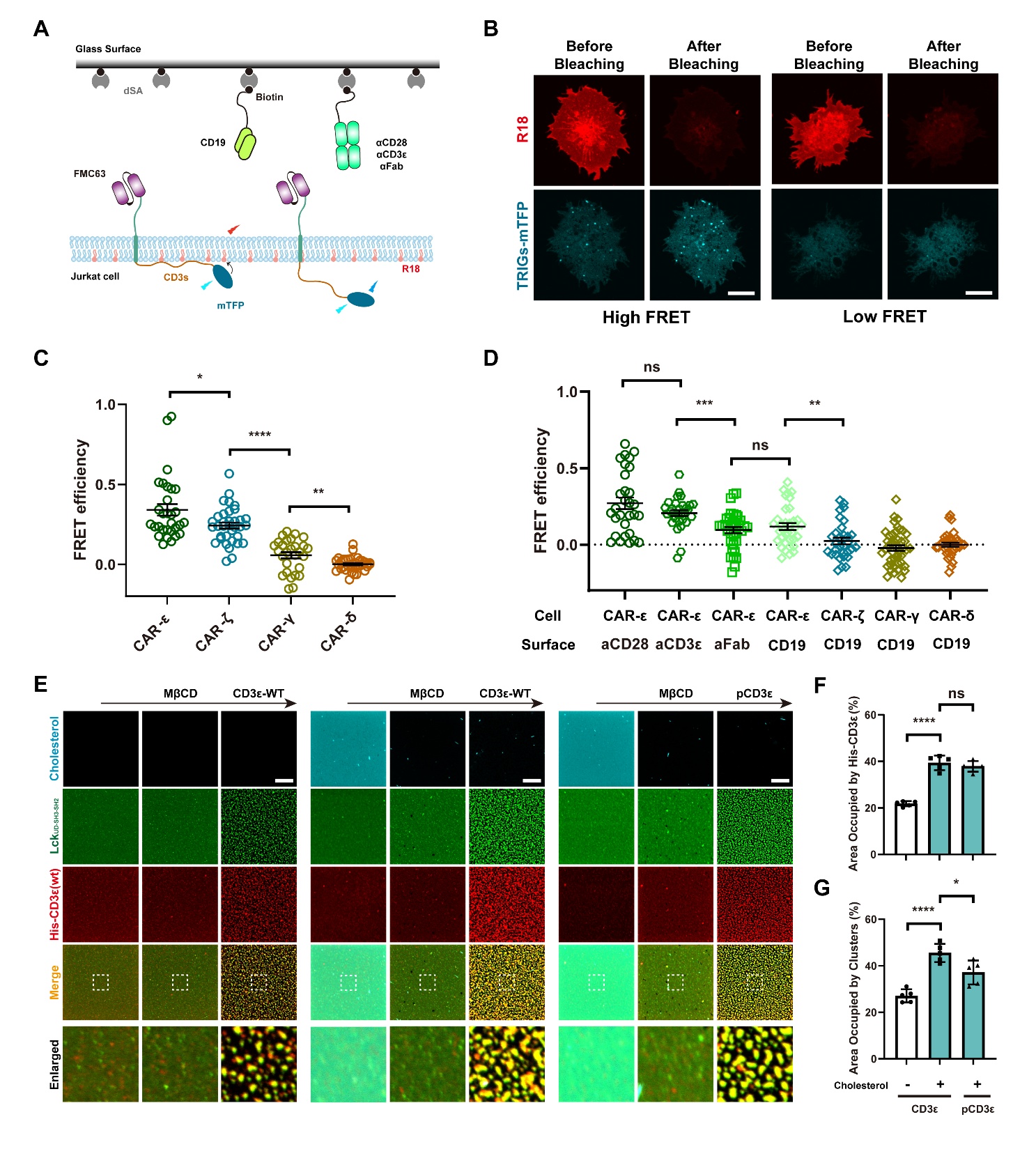

**Figure S1. State transitions of the CD3-ICD are governed by extracellular ligand binding and membrane tension.**

**A-B**. Schematic of CAR molecule design was shown in **A**, typical images of High FRET and Low FRET were shown in **B**, scale bar: 10 μm;

**C-D**. FRET efficiency between mTFP and membrane dye R18 measured by dequenching in live TRIG engineered Jurkat T cells, Jurkat cells were stimulated with immobilized antibodies as indicated (**C**: anti-β2m, **D**: anti-CD28, anti-CD3ε，anti-Fab and CD19 protein, data were shown as scattered plot and Mean±SEM, n>30; Unpaired t test; ns, *P* >0.05; *, *P* < 0.05; **, *P* < 0.01; ***, *P* < 0.001; ****, *P* < 0.0001);

**E-G**. Phase separation of Lck_UD-SH3-SH2_ with membrane-bound CD3ε in the presence of MβCD and CD3ε on SLB, the typical images were shown in **E** (scale bar: 10 μm**)**, the percentage of area occupied with membrane-bound CD3ε (**F**) and Lck_UD-SH3-SH2_/CD3ε clusters (**G**) were calculated, data were shown as Mean±SD (n=5; Unpaired t test; ns, *P* > 0.05; *, *P* < 0.05; ****, *P* < 0.0001).

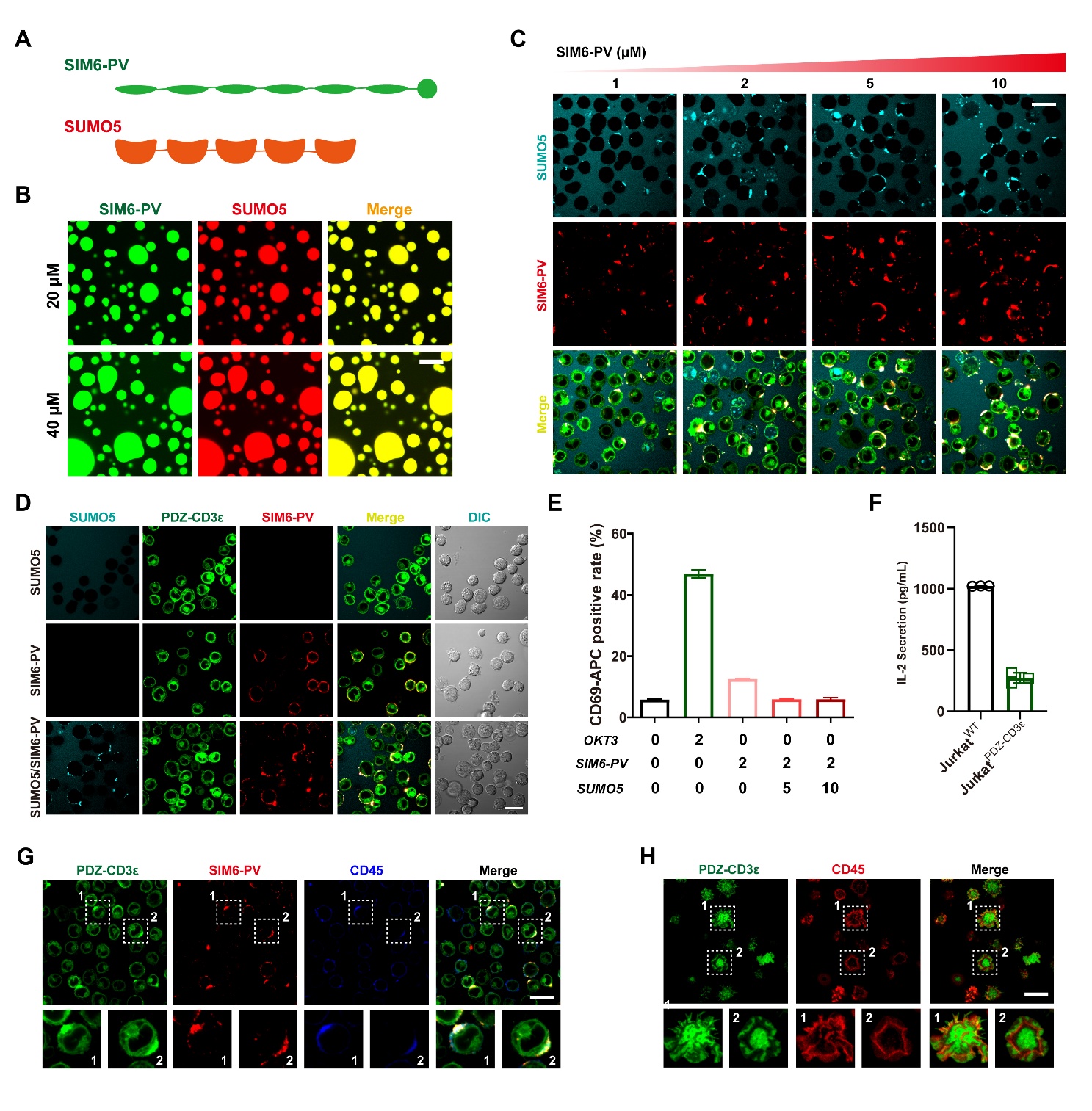

**Figure S2. Extracellular-domain phase separation–induced CD3ε clustering is insufficient to mediate T cell activation.**

**A**. Schematic of SIM6-PV and SUMO5 molecules;

**B**. Typical images of SIM6-PV/SUMO5 phase separated droplets, scale bar: 20 μm;

**C-D**. SIM6-PV/SUMO5 phase separation on the surface of PDZ-CD3ε engineered Jurkat cells, titration of SIM6-PV was shown in **C**, the distribution of PDZ-CD3ε in the presence of SUMO5, SIM6-PV and SIM6-PV/SUMO5 were shown in **D**, scale bar: 20 μm;

**E**. CD69 expression of PDZ-CD3ε engineered Jurkat cells stimulated with OKT3, SIM6-PV, SUMO5 and SIM6-PV/SUMO5, data were shown as Mean±SD, n=3;

**F**. IL-2 secretion of wild-type and PDZ-CD3ε engineered Jurkat cells stimulated with OKT3, data were shown as Mean±SD, n=3;

**G-H**. Typical images PDZ-CD3ε engineered Jurkat cells stimulated with SIM6-PV/SUMO5 (**G**) and membrane bound SIM6-PV (**H**), scale bar: 20 μm.

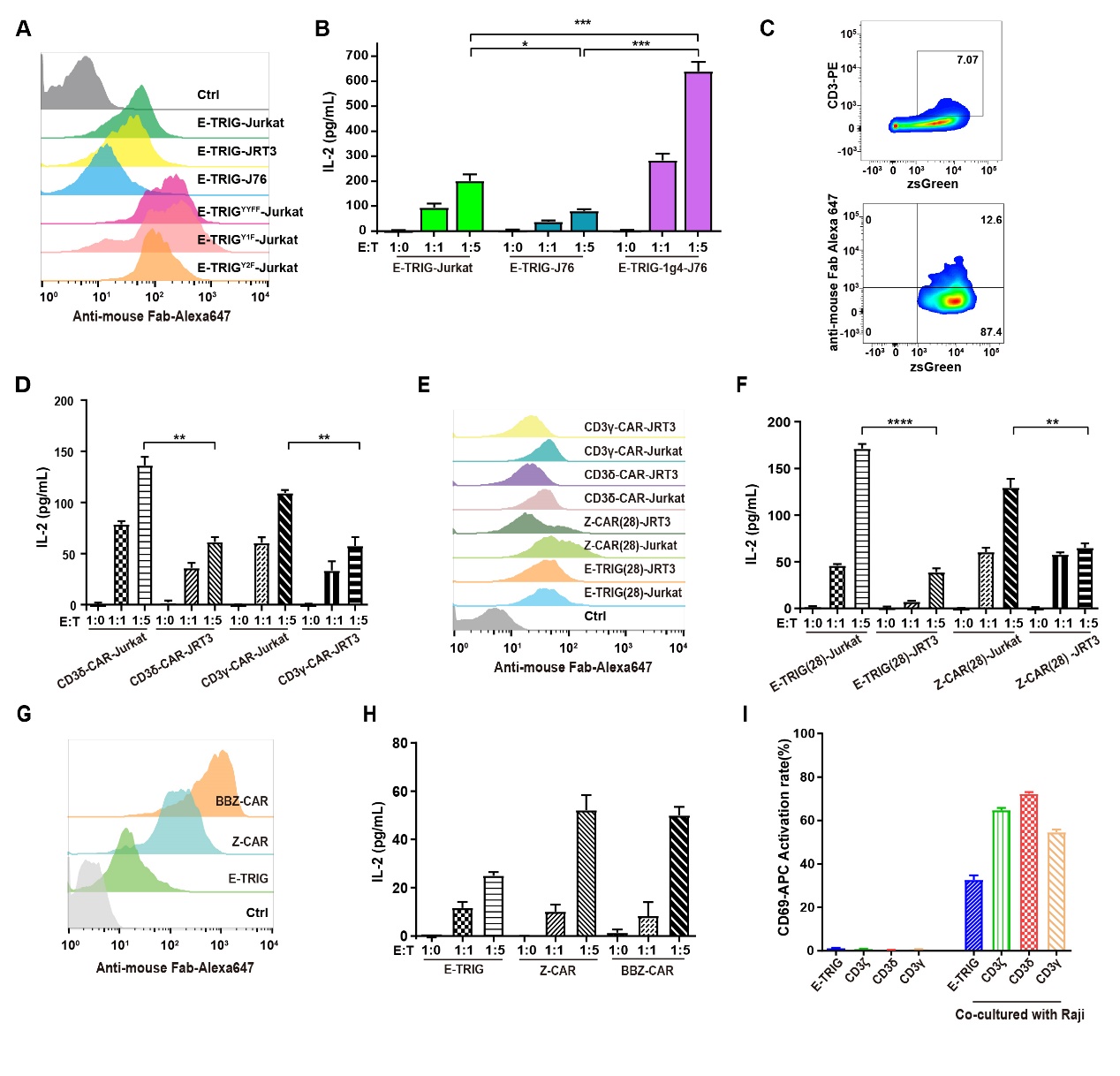

**Figure S3. T cell activation mediated by CD3ε intracellular domain-based TRIG requires endogenous TCR expression.**

**A.** Flow cytometry detection of membrane surface expression of various TRIGs on Jurkat cells;

**B.** ELISA analysis of E-TRIG-T, E-TRIG-J76 and E-TRIG-1g4-J76 cells cocultured with Raji B cells engineered with A2-MHC (n=3; Unpaired t test; *, *P* < 0.05; ***, *P* < 0.001);

**C.** Membrane surface expression of CD3 (1g4TCR successfully organized) and TRIG on J76 cells used in **B**;

**D.** IL-2 secretion of CD3 subunits-based CAR designs in Jurkat cells or JRT3 cells (n=3; Unpaired t test; **, *P* < 0.01);

**E.** Flow cytometry detection of membrane surface expression of various CARs on Jurkat cells or JRT3 cells used in **D and F**;

**F.** IL-2 secretion of TRIGs (CD28 transmembrane domain) in Jurkat cells cocultured with Raji B cells at different E:T ratios (n=3; Unpaired t test; **, *P* < 0.01; ****, *P* < 0.0001);

**G.** Flow cytometry detection of membrane surface expression of various CARs (anti-BCMA scFv) on Jurkat cells;

**H.** IL-2 secretion of CARs (anti-BCMA scFv) in Jurkat cells cocultured with Raji B cells at different E:T ratios (n=3);

**I.** CD69 activation of different CD3-subunits based CAR-Jurkat cells detected by flow cytometry.

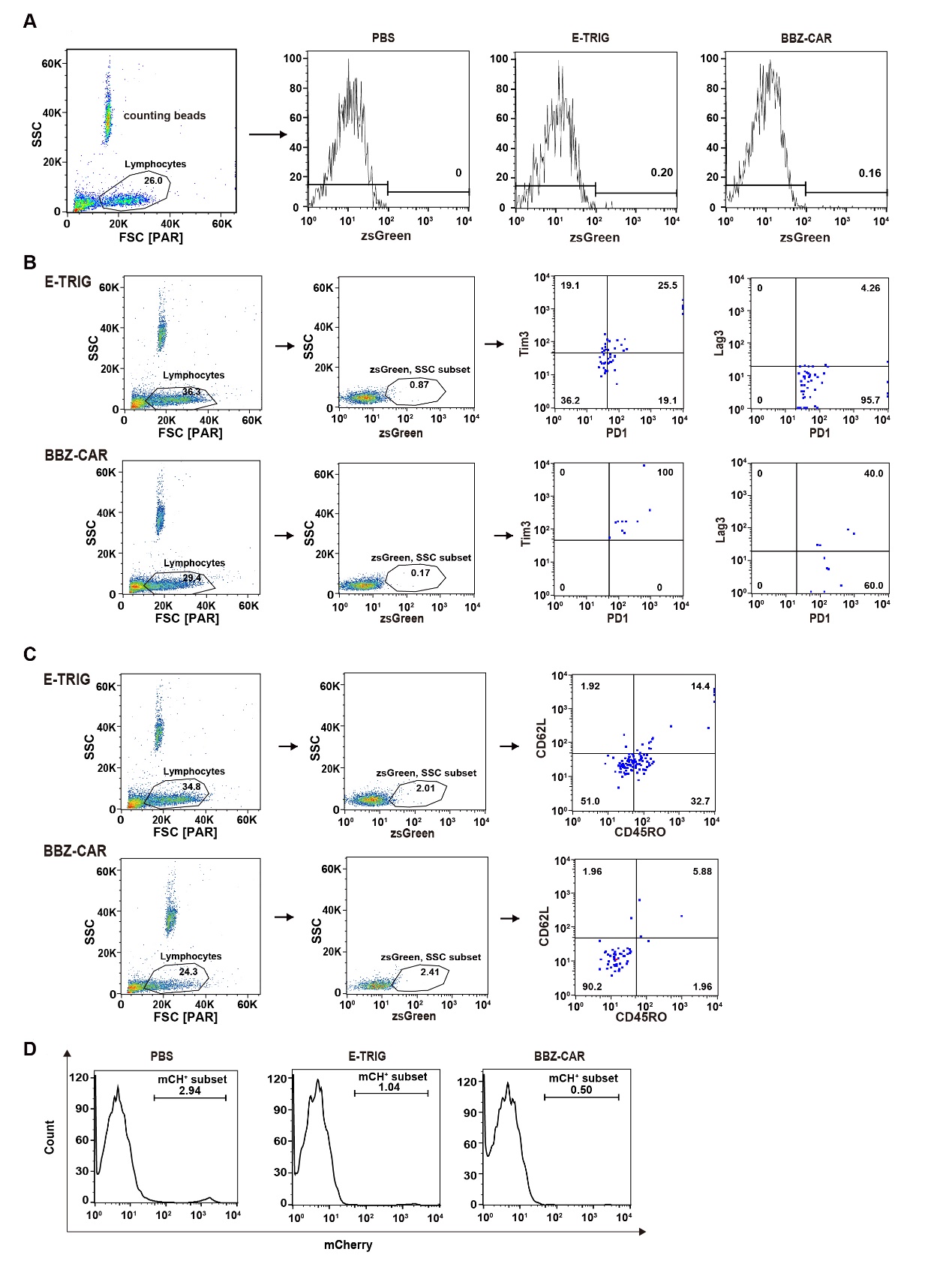

**Figure S4. TRIG-T cells exhibit a proliferative, memory-retaining, and exhaustion-reduced phenotype in vivo.**

**A.** Representative flow cytometry analysis for Figure 4E showing CAR-Ts persistence in vivo;

**B.** Representative flow cytometry analysis for Figure 4F indicating the exhaustion state of CAR-Ts in vivo;

**C.** Representative flow cytometry analysis for Figure 4G showing the CD45RO^+^CD62L^+^ CAR-T cells percentage;

**D.** Representative flow cytometry analysis for Figure 4H showing tumor burden percentage.

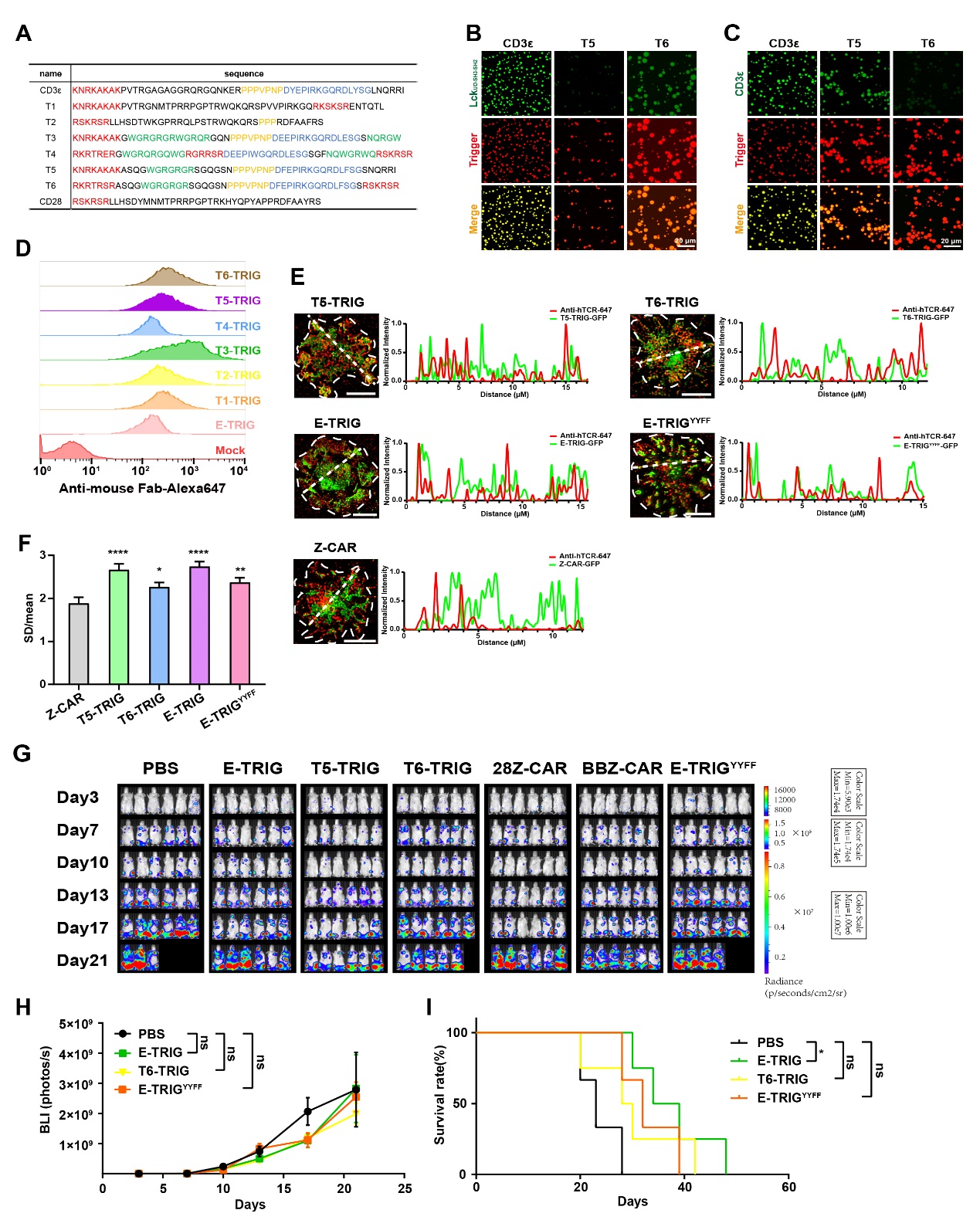

**Figure S5. In silico design of TRIG molecules for T cell engineering.**

**A**. Amino acids sequence and motif analysis of CD3ε, TRIG and CD28 intracellular domain, basic amino acid-rich regions (BRS) are marked in red, proline-rich regions (PRS) in yellow, ITAM or ITAM mutants in blue, and clustering motifs in green;

**B-C**. Typical images of Lck (**B**) and CD3ε (**C**) phase separated droplets induced by TRIGs intracellular domain, scale bar: 20 μm;

**D**. Membrane expression of TRIGs on primary T cells detected by flow cytometry;

**E**. Co-localization analysis of endogenous TCR with TRIGs and Z-CAR. TRIG engineered Jurkat cells were stimulated with SLB-immobilized CD19, cells were fixed and stained with APC ligated TCR antibody, Scale bar: 5 μm;

**F**. Quantification of TRIGs clustering in Jurkat cells shown by SD/mean. TRIG engineered Jurkat cells were stimulated with slide-immobilized CD19 and fixed. Significance comparisons are shown for all groups against the Z-CAR group, (n=20; Unpaired t test; *, *P* < 0.05; **, *P* < 0.01; ****, *P* < 0.0001);

**G**. Representative bioluminescence imaging of mouse model after CAR-Ts treatment. For all groups, n=6;

**H**. Quantification of PBS, E-TRIG, T6-TRIG and E-TRIG^YYFF^ groups in Figure S5G showing the tumor burden as indicated by total radiance (p/s). Two-way ANOVA with Tukey’s multiple-comparisons test was used to assess significance, (n=6; ns, *P* > 0.05);

**I**. Survival curve of PBS, E-TRIG, T6-TRIG and E-TRIG^YYFF^ groups in Figure S5G. Mantel–Cox test was used to assess significance, (n=6; Unpaired t test; ns, *P* > 0.05; *, *P* < 0.05).

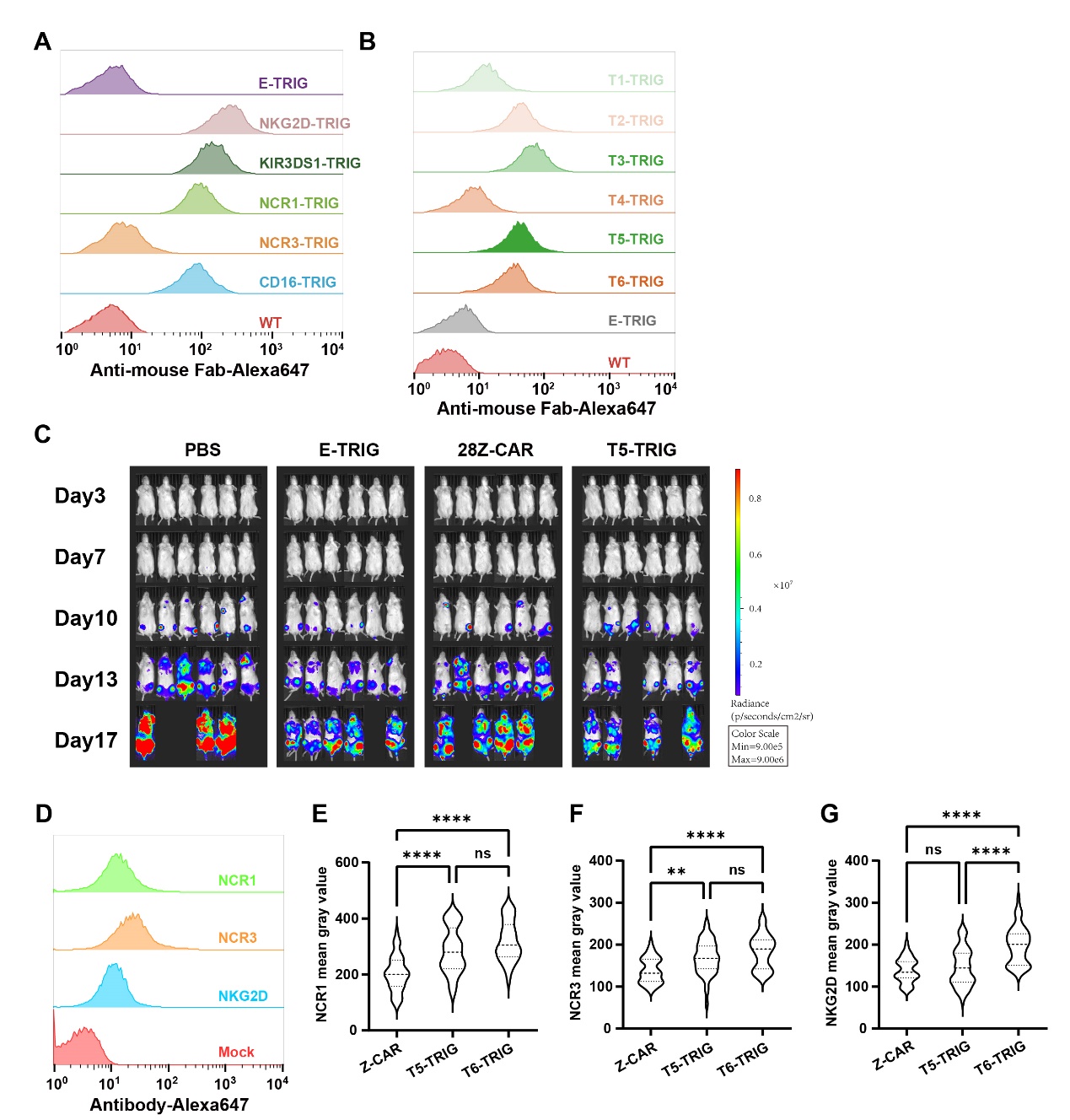

**Figure S6. Harnessing phase separation-based trigger mechanism endows NK cell liquid tumor killing ability.**

**A-B**. Membrane expression of NK receptor-TRIGs (A) and T1-T6 TRIGs (B) on NK92MI cells detected by flow cytometry;

**C**. Representative bioluminescence imaging of mouse model after TRIG-NK treatment. For all groups, n=6;

**D**. Membrane expression of NCR1, NCR3 and NKG2D on NK92MI cells detected by flow cytometry;

**E-G**. Quantification of TRIG-induced clustering of endogenous receptors in NK cells from TIRF-SIM fluorescence imaging, represented by the mean gray value of NCR1 (**E**), NCR3 (**F**) and NKG2D (**G**). TRIG engineered NK92MI cells were stimulated with SLB-immobilized CD19, cells were fixed and stained with APC ligated NCR1, NCR3 or NKG2D antibody (n=50 for all three groups; Unpaired t test; ns, *P* > 0.05; **, *P* < 0.01; ****, *P* < 0.0001).
